# Global loss of promoter-enhancer connectivity and rebalancing of gene expression during early colorectal cancer carcinogenesis

**DOI:** 10.1101/2022.08.26.505505

**Authors:** Yizhou Zhu, Hayan Lee, Shannon White, Annika K. Weimer, Emma Monte, Aaron Horning, Stephanie A. Nevins, Edward D. Esplin, Kristina Paul, Gat Krieger, Zohar Shipony, Roxanne Chiu, Rozelle Laquindanum, Thomas V. Karathanos, Melissa WY Chua, Meredith Mills, Uri Ladabaum, Teri Longacre, Jeanne Shen, Ariel Jaimovich, Doron Lipson, Anshul Kundaje, William J. Greenleaf, Christina Curtis, James M. Ford, Michael P. Snyder

## Abstract

Although 3D genome architecture can be essential for gene regulation, the biological implications of long-range chromatin interactions in disease remain elusive. In this study, we traced the early evolution and malignant transformation of colorectal cancer by generating high-resolution chromatin conformation maps of 33 colon samples spanning different stages of early neoplastic growth from polyps of Familial Adenomatous Polyposis (FAP) patients. Our analysis reveals a substantial progressive loss of genome-wide cis-regulatory connectivity at early stages of malignancy, which correlates with a non-linear effect on gene regulation. Genes with high promoter-enhancer (P-E) connectivity in unaffected mucosa are not correlated with elevated baseline expression, but instead tend to be up-regulated at advanced stages. Inhibition of highly connected promoters preferentially represses gene expression in colorectal cancer cells relative to normal colonic epithelial cells. Our results suggest a two-phase model whereby neoplastic transformation reduces P-E connectivity from a redundant state to a rate-limiting one for transcriptional levels. Overall, our study illuminates the intricate interplay between 3D genome architecture and gene regulation during early colorectal cancer progression, and provides valuable insights for potential therapeutic interventions targeting the connectivity of cis-regulatory elements.

## Introduction

The technological advent of 3D chromosome organization mapping has revealed important insights into genome folding^1–3^. Multilayered structures maintained by molecular contacts, insulators, and aggregative domains together compact the 2-meter DNA into non-random spatial configurations in the nucleus^2,4^. However, the functional implications of such spatial organization on fundamental biological processes remain largely elusive.

A key discovery in genome topology is the formation of topologically associating domains (TADs)^5,6^, high-order structures that partition the genome into contiguous regions through a proposed loop extrusion mechanism^7,8^. While genetic mutations affecting TAD structures have been linked to oncogenic gene dysregulations in specific cases^9–11^, the exact role of TAD organization in transcription regulation remains an open question. Recent studies have shown a surprisingly moderate transcriptional response to the manipulation of boundary elements such as CTCF and cohesin components^12,13^. Furthermore, computational approaches have suggested a lack of co-expression between genes residing in the same TAD^14,15^. These findings imply that gene regulation is often not particularly constrained by large sub-megabase folding domains, but rather depends on a finer layer of regulatory architecture at the sub-TAD level.

Recent advancements in chromatin conformation capture technologies, utilizing micrococcal nuclease^16,17^ or a combination of restriction enzymes^18,19^, have improved mapping resolution and enhanced the detection of sub-TAD structures. These methodologies have uncovered prevalent distal interaction activities, including architectural stripes and insulation activities associated with active cis-regulatory elements. However, the functional implications of these structures remain largely uncharacterized. Moreover, studies of transcriptional kinetics based on imaging^20,21^ and multi-omics sequencing^22,23^ have revealed disjoined changes between the spatial proximity of regulatory elements and transcriptional activation events. These observations highlight the intricate role of physical connectivity in gene regulation.

Colorectal cancer (CRC) represents a major global health burden, and is the second leading cause of cancer death in the United States^24^. Over 80% of colorectal carcinomas are initiated by loss-of-function mutations of APC, a key component in the cytosolic complex that targets β-catenin for destruction and suppresses Wnt signaling^25,26^. Familial Adenomatous Polyposis (FAP) patients carry germline APC mutations and develop tens to thousands of precancerous polyps at different stages and sizes as well as occasional adenocarcinomas; these polyps are believed to represent early stages of CRC^27,28^. Thus, the study of polyps at different stages of development in FAP patients provides a valuable model for studying the cascades of epimutations and gene dysregulations during early oncogenesis.

In the context of the Human Tumor Atlas Network (HTAN)^29^, we profiled genome-wide chromatin conformation at up to 100 bp resolution in 33 colon samples from FAP and CRC patients representing different stages of CRC progression. By integrating these data with transcriptome and epigenome profiling, we investigated the relationship between fine-gauge chromatin structures organized by active regulatory elements and gene dysregulation associated with polyp malignancy. Our analysis revealed a progressive loss of cis-regulatory connectivity from mucosa to polyps to adenocarcinoma, corresponding with dysregulated gene expression in a non-linear fashion. The initial connectivity levels prior to oncogenic progression may be a key factor in this process. We propose that the remodeling of promoter-enhancer (P-E) connectome is not only indicative of alterations of gene expression, but also reflects shifts in the transcriptional response to epigenetic alterations in polyps and adenocarcinoma. Our rich dataset provides a valuable resource for unraveling the chromatin architectural basis of early CRC development.

## Results

### Fine-mapping distal interactions of active regulatory elements using multi-digested Hi-C

To examine the chromatin architecture associated with regulatory elements in clinical tissue samples, we developed multi-digested Hi-C (mHi-C), a protocol derived from in situ Hi-C^30^ that utilizes five 4-cutter restriction enzymes and moderated detergent conditions to achieve ultrafine mapping (mean fragment size = 52 bp) of the distal chromatin interactions^18,31^ (Fig **1A**, **S1A, Methods**). We generated mHi-C data for 33 frozen colon tissue samples at different stages of neogenesis (Fig **1B**), comprising 7 non-neoplastic mucosa, 19 dysplastic polyps, and one adenocarcinoma from 4 familial adenomatous polyposis (FAP) patients, as well as six additional adenocarcinoma samples from non-FAP individuals who developed sporadic colorectal cancer (CRC). A total of 1.59 billion unique intrachromosomal long-range (≧ 1kb) interaction contacts were mapped (Fig **S1B,C**).

**Figure 1.**
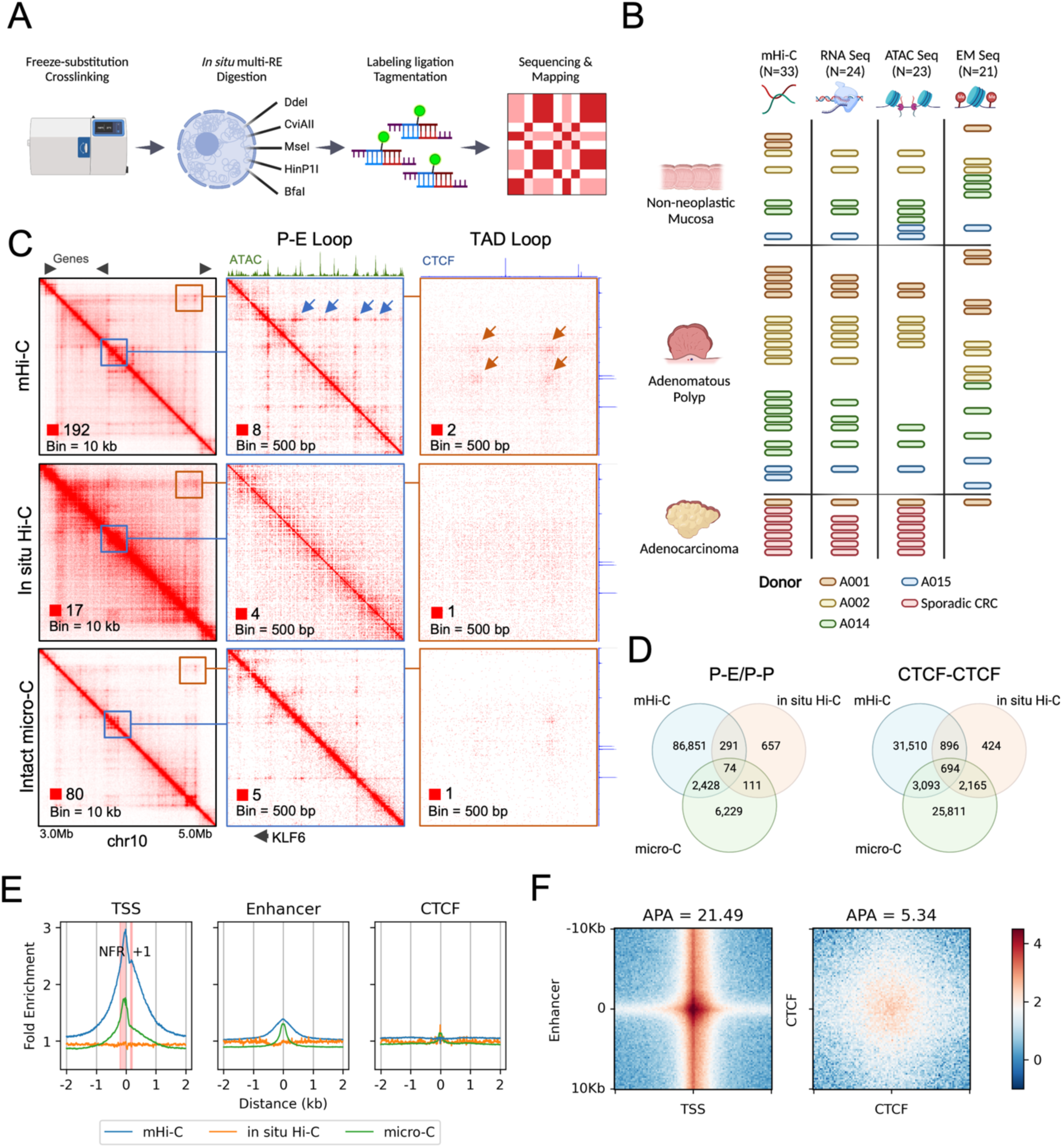
mHi-C reveals interactions associated with active cis-regulatory elements. (A) Schematic representation of the mHi-C workflow (B) Summary of colon tissue samples analyzed by multi-omics assays. Bar colors represent different donors. Each row corresponds to a unique donor patient with sporadic colorectal cancer. (C) Comparison of contact matrices generated from mHi-C (combined colon tissues) with in situ Hi-C (HCT116) and intact micro-C (HCT116) at various resolutions. Blue arrows highlight interaction dots formed between the KLF6 promoter and adjacent enhancers. Orange arrows show structural loops at TAD boundaries. (D) Venn diagrams illustrating the overlap of interaction loops identified by the three methods. (E) Average fold enrichment of distal interactions at transcription start sites (TSS), active enhancers, and CTCF binding sites in mucosa samples. Red intervals indicate the nucleosome-free region (NFR) and the +1 nucleosome regions upstream and downstream of the TSS, respectively. (F) Aggregated peak analysis (APA) of loops between promoters and active enhancers (N=9,174) and between CTCF-CTCF (N=30,208).

Similar to micro-C^16,17,32^, mHi-C robustly revealed fine-gauge structures at sub-kilobase resolution (200 bp – 1 kb). This included dot interactions, indicative of looping of two fixed anchors, as well as architectural stripes, indicative of dynamic looping between a fixed anchor and the sliding intervening neighboring regions (Fig **1C**, **S2**)^7,8^. Previous studies have correlated these structures with interaction hotspots identified by high-depth 4C assays, enriched at regulatory elements^18,33,34^. To annotate the interacting regions, we mapped open chromatin regions in 23 matched samples using ATAC-Seq^35^ (Fig **1B**), and annotated the regulatory elements by using the Ensembl regulatory build^36^. Notably, both micro-C and mHi-C, but not in situ Hi-C, revealed enriched interactions at promoters and enhancers (Fig **S1D**). This enrichment was specific to long-range contacts, which persisted after normalizing against short-range or total read coverage (Fig **1D**, **S1E**), indicating that the observed stripe signals are not artifacts of differential contact mappability due to the high accessibility of these regions.

Using the HICCUPs algorithm^37^, we identified 279,480 loop interactions, including 91,706 promoter-promoter (P-P) or promoter-enhancer (P-E) contacts (Fig **S1F**). Compared to chromatin conformation profiles obtained from intact micro-C and in situ Hi-C, mHi-C identified approximately 10 and 100 fold more P-E/P-P interactions, but only 1.5 and 8.5 fold more loops between CTCF binding sites, respectively (Fig **1E**). This suggests that mHi-C specifically improves the detectability of contacts among active regulatory elements. Meanwhile, using a peak calling algorithm based on MACS2^18,38^, we identified 254,642 stripes in all samples (see Methods). Of the 279,480 loops, 266,419 (95%) overlapped with stripes (Fig **S1G**), indicating that the majority of loop contacts are formed along with stripe extension.

At 100 bp resolution, we observed a significant difference between promoter-enhancer and CTCF-CTCF interactions. Whereas contacts between promoters and enhancers are formed at open chromatin regions, CTCF interactions are not strictly overlapping with CTCF binding sites. Instead, they spread widely within a multi-kb flank region (Fig **1F**). These patterns are consistent with occupancy of the cohesin components at the open chromatin of active regulatory elements, as opposed to being retained at a broader region around CTCF sites (Fig **S1H**). Furthermore, we found that the loop signals of P-E interactions were comparable with the sum of the anchors’ stripe strengths, whereas the loop strengths of CTCF interactions significantly exceeded their architectural stripe strengths (Fig **2A**, **S3A**). This difference suggests that, unlike CTCF boundaries, which can maintain stable loop structures^39^, promoter-enhancer interactions are dynamically maintained while the intervening anchors interact frequently with each other’s neighborhoods.

**Figure 2.**
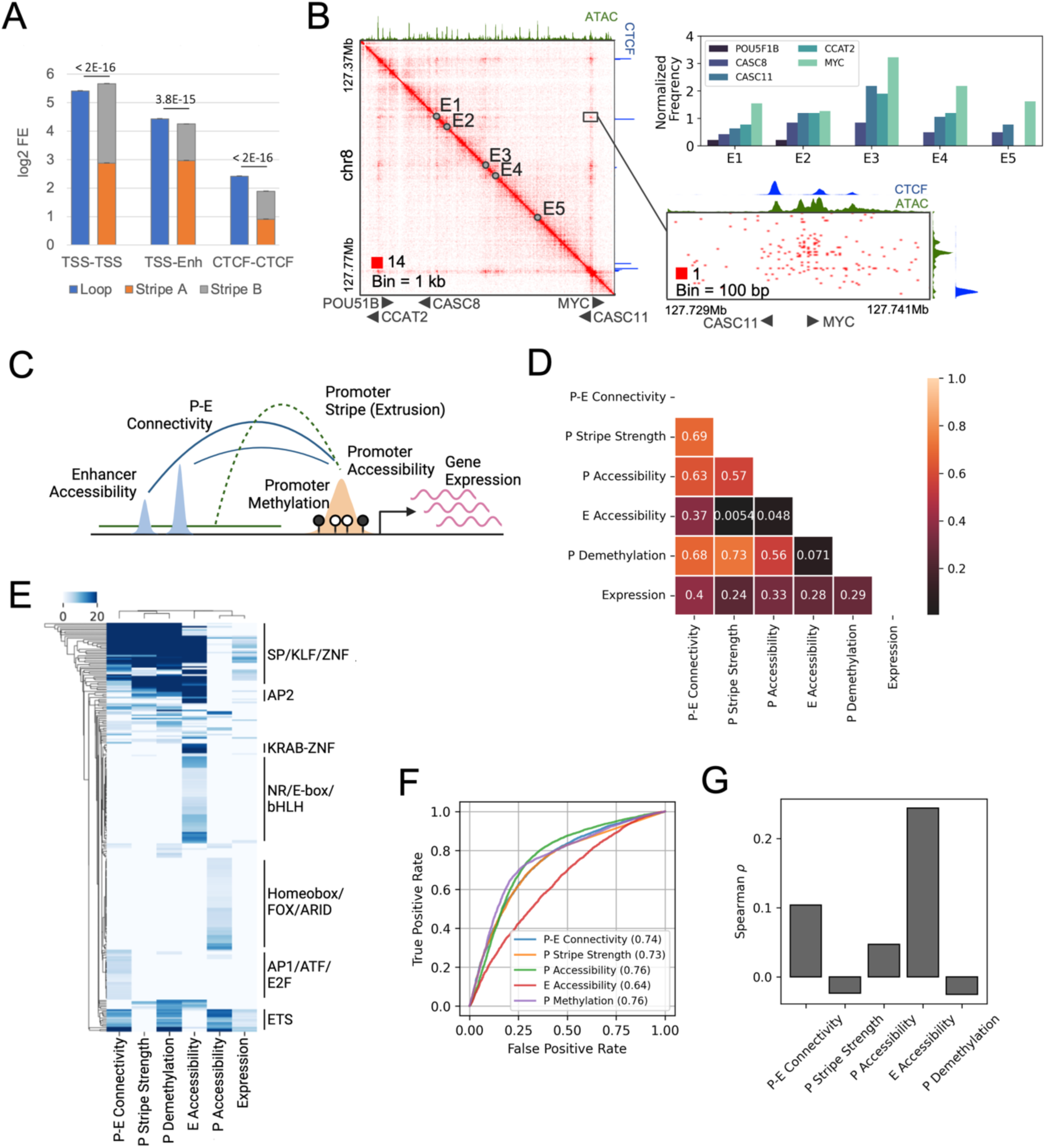
Correlation between promoter stripe formation and P-E connectivity. (A) Comparison of loop strengths with the logarithmic sum of stripe strengths at the anchors for loops formed between different regulatory elements. Error bars represent confidence intervals. Statistical significance was assessed using the Wilcoxon signed-rank test. (B) Contact heatmap (combined colon tissues) of the MYC upstream cancer risk locus. The top right panel shows contact frequencies of resident genes with the five putative enhancers exhibiting the highest distal interaction activity. A detailed view of contact distribution between enhancer E1 and the MYC/CASC11 genes is provided in the bottom right panel. (C) Schematic representation of the integration of conformational, epigenetic, and transcriptional features for downstream analysis. (D) Spearman correlation matrix of average feature strengths in mucosa samples for all examined coding genes (N=14,692). (E) Hierarchical clustering of genes exhibiting top 10% intensity for any of the six examined features, based on the degree of motif enrichment (adjusted −log10(p-value)) on their promoters. (F) Receiver Operating Characteristic (ROC) analysis for predicting actively expressed genes (transcripts per million (TPM) > 0.5, N=10,663) using various structural and epigenetic features. Numbers indicate the area under the curve (AUC) scores. (G) Spearman correlation of the expression levels of actively expressed genes with the examined features.

### Architectural stripes formed by promoters shape gene-specific P-E connectivity

We observed that promoter-enhancer interactions are asymmetrically contributed by the relatively stronger stripe-forming activity of the promoters and weaker activity of the enhancers (Fig **1F**, **2A**). This asymmetry underscores the dominant role of promoters in shaping the promoter-enhancer (P-E) connectivity. To validate this hypothesis, we conducted a case study of the MYC upstream locus, which is a known risk hotspot for multiple cancer types^40^ that resides near five coding genes (POU51B, CCAT2, CASC8, CASC11, and MYC) and multiple enhancers. Our results demonstrated that MYC, which exhibited the highest stripe strength, consistently interacted with all enhancers with the highest frequency among all five genes, despite other genes being located closer to these enhancers (Fig **2B**). Remarkably, the enhancers tended to bypass CASC11, a gene located only 1 kb upstream of MYC with substantial promoter CTCF binding, and instead favored robust interactions with the MYC promoter, which showed lower CTCF affinity. We extended our examination to additional loci and consistently found that gene-specific promoter stripe activities lead to distinctive interaction profiles for genes sharing the same enhancer context (Fig **S3B**).

To further investigate the relationship between P-E connectivity and promoter/enhancer activity, we profiled the landscapes of methylome and transcriptome from the matching FAP/CRC samples (Fig **1B**). We quantified the connectivity of all coding genes with their neighboring enhancers within 200 kb distance. We then correlated this connectivity with the accessibility of the enhancers, the methylation, and stripe activity of the gene promoters, assessed by the fold enrichment of the overlapping architectural stripes, in unaffected mucosa (Fig **2C**). The connectivity exhibited a strong correlation with stripe strength (*ρ*=0.69), demethylation (*ρ*=0.68), and accessibility (*ρ*=0.63) of the gene promoter. By contrast, its correlation with enhancer accessibility was much lower (*ρ*=0.37) (Fig **2D**). Consistently, clustering analysis revealed distinct groups of genes associated with high connectivity but low-to-moderate enhancer accessibility, and vice versa, indicating a significant discrepancy between the availability of enhancers and their connectivity with promoters (Fig **S3C**). Motif analysis uncovered a strong enrichment of transcription factors (TFs) with GC-rich motifs on highly interactive promoters, distinguishing them from promoters associated with high accessibility or rich neighboring enhancer contexts (Fig **2E**). Collectively, these results suggest that the degree of P-E connectivity of genes is predominantly correlated with the stripe activity on their promoters, corroborating our observations in the case studies.

Upon observing a significant divergence between the promoter’s connectivity and the availability of distal regulatory elements, we investigated which factor was more relevant to transcription. We discovered that the overall correlation of connectivity with gene expression surpassed that of enhancer accessibility (Fig **2D**). However, this high correlation predominantly stemmed from the association of transcriptionally inactive genes (TPM <0.5) with low connectivity (Fig **S3D**). As such, high connectivity serves as a robust indicator of an “on” state for promoters, similar to their accessibility, demethylation, and stripe strength (Fig **2F**). Conversely, the expression level of active genes (TPM >0.5) was poorly correlated with the connectivity (*ρ*=0.10) but rather explained by the enhancer accessibility (*ρ*=0.24) (Fig **2G**). This implies that regulation of quantitative expression was dependent on the activity of enhancer context for genes with connectivity passing beyond the “on” threshold.

### Impaired stripe formation and connectivity indicates promoter instability in early colon cancer development

During the development of CRC malignancy, we observed that more than 80% of identified stripes and loops in polyps and over 90% in cancer showed reduced signals (Fig **3A**, **S4A**). Among the annotated CREs, gene coding transcription start sites (TSSs) exhibited an exceptionally high loss rate of stripe strength (Fig **S4B**). Consistently, the global P-E connectivity was progressively lost in advanced stages, suggesting reduced promoter-enhancer communications associated with stripe loss (Fig **3B, C**). To test whether these alterations are due to increased chromosomal rearrangements along with stage progression, we applied EagleC^41^ to the mHi-C results, identifying 1 to 17 structural variants (SVs) in each sample (Fig **S4C**). The sparsity of the SVs and a comparable number of average events between the mucosa (2.5) and polyp (2.5) stages suggests that they are unlikely a primary driving factor for the genome-wide loss of interactions.

**Figure 3.**
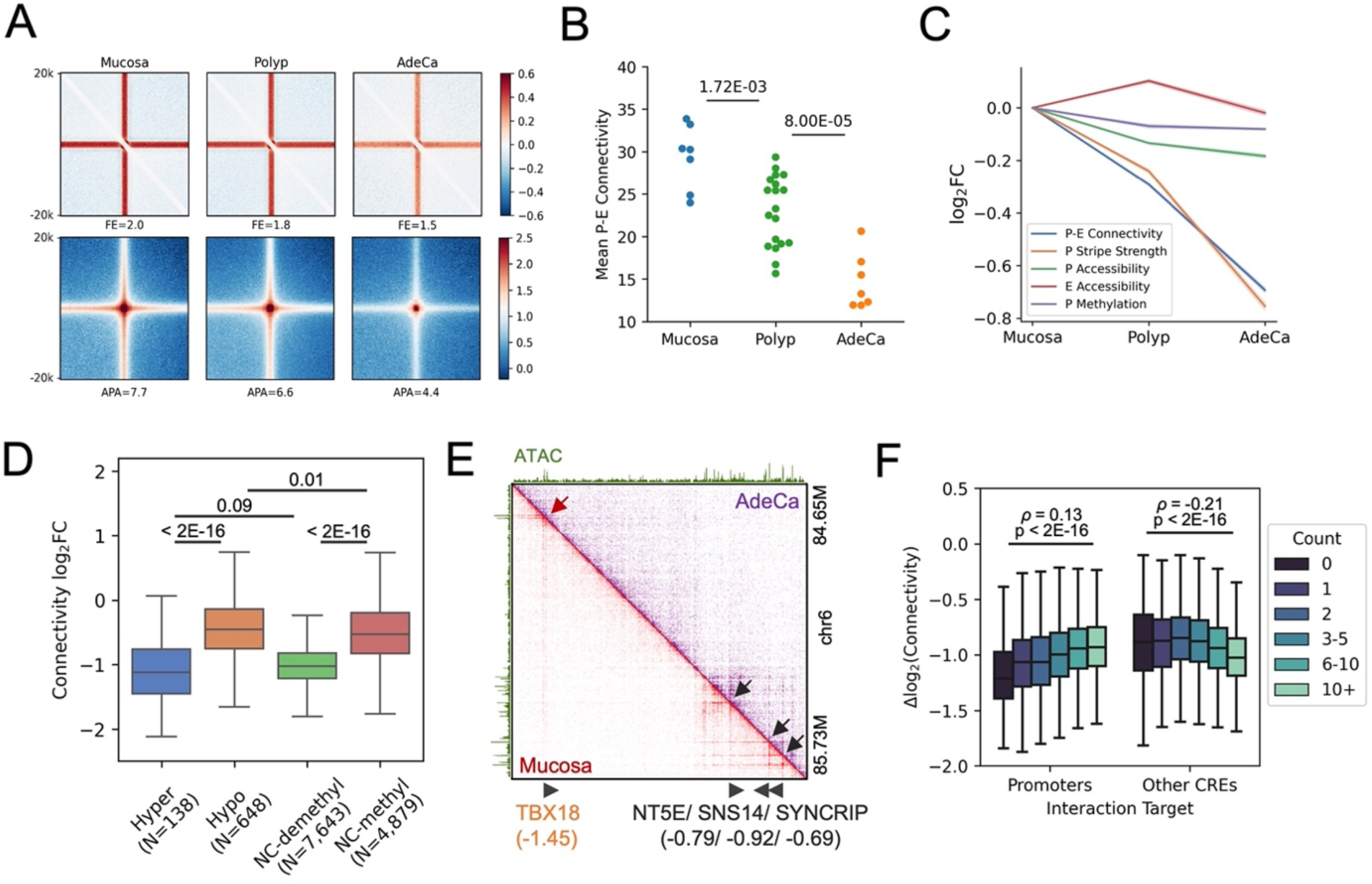
Loss of distal connectivity in polyps and adenocarcinoma. (A) Aggregated peak analysis (APA) of all stripe (top) and loop (bottom) anchors, with fold enrichment of signals indicated below the panels. (B) Mean promoter-enhancer (P-E) connectivity for coding genes in mucosa, polyp, and adenocarcinoma (AdeCa) samples, with p-values from the Mann-Whitney U test presented above. (C) Average log2 fold changes of structural and epigenetic features in polyps and adenocarcinoma, with confidence intervals represented by shaded areas. (D) Changes in connectivity between mucosa and adenocarcinoma for genes categorized by hypermethylation, hypomethylation, or no change (NC, <5% difference) in methylation status. Groups with demethylated (<25%) and methylated (>40%) promoters are compared. (E) Comparison of contact heatmaps for a representative locus in mucosa and adenocarcinoma samples, with log2 connectivity changes for gene promoters indicated below. (F) Comparison of connectivity changes for genes interacting with varying numbers of promoters and other cis-regulatory elements (CREs), including Spearman correlation coefficients and p-values from the Mann– Whitney U test.

To investigate whether the loss of P-E connectivity was due to changes in the activity of the regulatory elements, we aligned the results with the chromatin accessibility and methylation profiles. In both polyps and adenocarcinoma, alterations in connectivity were poorly correlated with the accessibility changes of both gene promoters and neighboring enhancers (*ρ*≤0.13) (Fig **S4D**). We also observed a mild progressive loss of accessibility on the promoters but not on the enhancers (Fig **3C**). However, the degree of accessibility loss (−12.3%) was marginal compared to the substantial losses of P-E connectivity (−39.3%) and promoter stripe strength (−41.8%), suggesting underlying factors that specifically contributed to the impairment of distal interaction. On the other hand, hyper- and hypo-methylated promoters were associated with high- and low-connectivity losses, respectively (Fig **3D**), consistent with the well-characterized repressive function of DNA methylation^42^. However, for the majority (>80%) of the promoters that were neither hypo- nor hyper-methylated, we found that demethylated promoters were also associated with a higher rate of connectivity loss compared to methylated ones (Fig **3D**, **S4E**). Furthermore, demethylated promoters that are hyper-methylated in the advanced stages are implicated by their significantly lower initial connectivity in the mucosa samples (Fig **S4F**). These results together indicate a common factor driving both the global connectivity loss of promoters and selective hyper-methylation of low connectivity ones, rather than hyper-methylation as a driving force of the connectivity loss.

Recent studies of clusters of enhancers, also known as super-enhancers, suggested that high valency and number of components in enhancer clusters increased their cooperativity through phase separation^43,44^. Inspired by this observation, we examined whether the stability of the P-E networks was affected by the valency of the networks. Interestingly, we found that a high valency of interacting promoters, but not enhancers, was associated with a lower rate of connectivity loss from mucosa to adenocarcinoma (**Fig 3E, F**). This suggests that cooperative promoter-promoter (P-P) interaction networks are associated with elevated stability during neogenic progression.

### Initial P-E connectome primes gene dysregulation during malignancy transition

To elucidate the implications of P-E connectivity on transcriptional outcomes, we analyzed 2,872 genes that exhibited progressive up- or down-regulation in polyps and adenocarcinomas (Fig **S5A, B**). We discovered that both up- and down-regulated genes displayed a similar degree of connectivity loss compared to the genome average (Fig **S5C**). Consistently, genes associated with increased or decreased connectivity loss did not correlate with up- or down-regulation (Fig **S5D**), indicating that the direct impact of P-E connectivity changes on differential gene expression was insignificant. Importantly, however, the first principal component of both the transcriptome and the P-E connectome revealed a consistent trajectory of stage progression, suggesting that the remodeling of the two “omes” during CRC development were closely related, despite their low linear correlation at each individual genes (Fig **S5E**).

Upon investigating the potential non-linear relationship between P-E connectivity and gene expression, we identified a significant correlation between differential gene expression and the levels of their connectivity relative to the genome average, as well as to their transcription levels (Fig **4A**). While up- and down-regulated genes were associated with high and low connectivity, respectively, the connectivity levels were established in unaffected mucosa samples rather than gained or lost in advance stages. As the stage progressed, the gene expression shifted towards higher correlation with the levels of P-E connectivity (Fig **4B**, **S6A**). Interestingly, a similar trend was also observed between gene expression and other indicators of promoter activity, such as accessibility, stripe activity, and demethylation, but not with the accessibility of neighboring enhancers (Fig **4B**). Collectively, these observations suggest a scenario in which impaired P-E connectivity in polyps and adenocarcinomas correlated with an increased dependence of transcription dosage control to the promoter activity.

**Figure 4.**
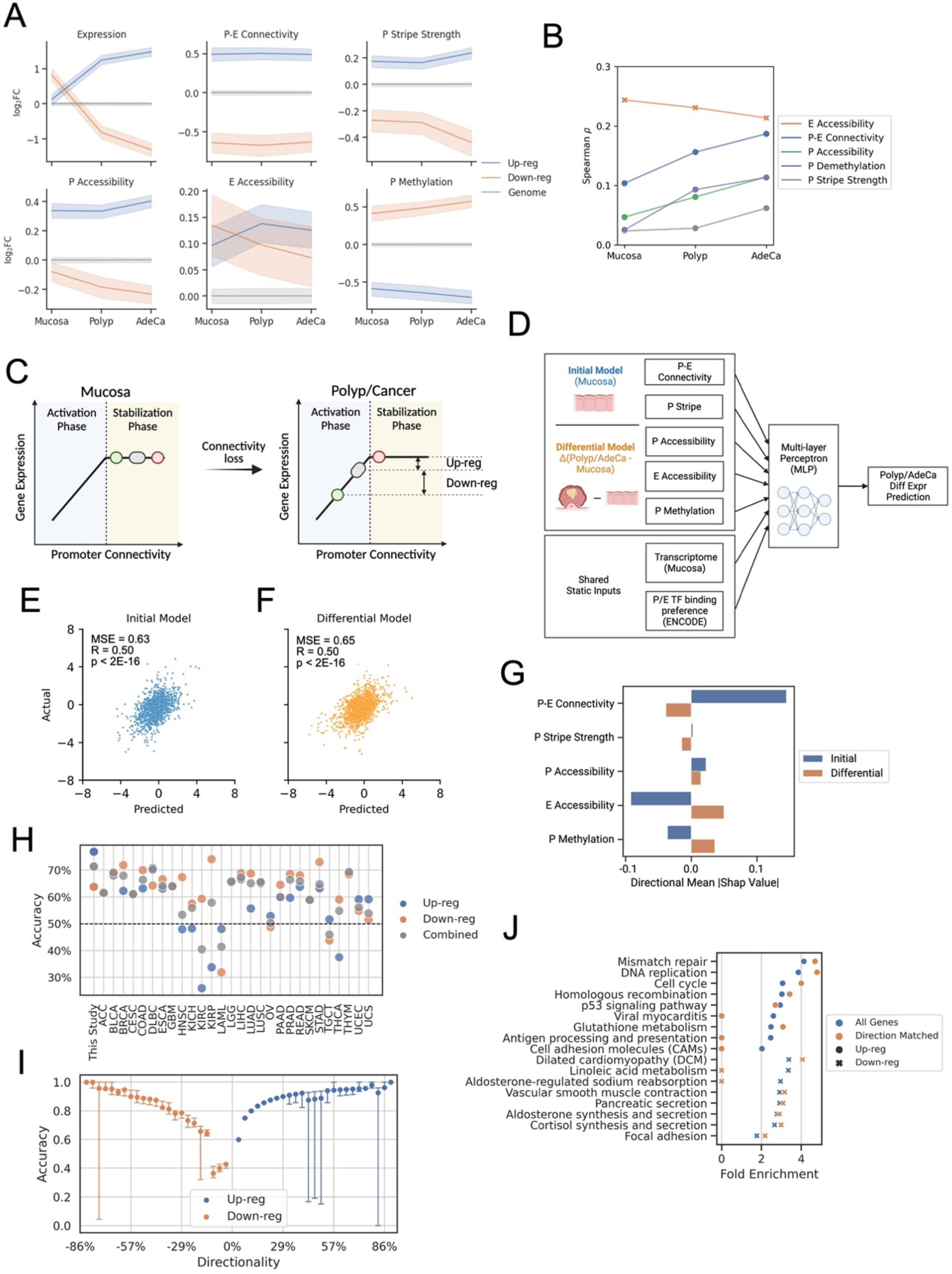
Predictable cancer gene dysregulation by initial P-E connectivity. (A) Relative fold changes of features for genes upregulated (N=1,089) or downregulated (N=944) in both polyps and adenocarcinoma, compared to the genome average at each stage, with confidence intervals shown as shaded areas. (B) Spearman correlations between transcription levels of active genes (TPM > 0.5) and their structural and epigenetic features at different stages of progression. (C) Schematics for the two-phase model. In normal conditions, most genes are in the saturated stabilization phase, where increased levels of P-E connectivity stabilizes the networks but do not contribute to higher gene expression. In polyp and cancer conditions, genes shift to the activation phase due to global losses of the connectivity, where expression levels are rate-limited by the connectivity levels. Alterations of gene expression during stage progression are therefore determined by their initial distance to the activation phase at normal condition. (D) Diagram illustrating the construction of “initial” and “differential” prediction models. (E) Predictive accuracy of the “initial” model for gene expression changes in polyps and (F) the “differential” model for a test set of genes (N=2,800). (G) The importance of features and the average direction of association of structural and epigenetic features in the predictive models. (H) Accuracy of the “initial” mucosa-polyp model in predicting the direction of significant expression changes in 28 cancer types from the TCGA database. (I) Prediction accuracy for genes grouped by their directionality scores. (J) Pathway ontology analysis for genes with altered expression in any TCGA cancer type versus those with accurately predicted directional changes by the “initial” model. Zero values indicate no significant enrichment (FDR > 0.1).

### A two-phase model of P-E connectivity predicts conserved transcription dysregulation in cancers

The significant association of cancer dysregulation with the P-E connectome prior to cancer development can be explained by a two-phase model. In unaffected mucosa, high P-E connectivity displays functional redundancy, which does not drive high gene expression but rather increases the stability of the P-E networks (stabilization phase). With the loss of connectivity during development of malignancy, the impaired connectivity of the promoters becomes a rate-limiting factor (activation phase), thereby establishing an increased linear correlation with gene expression (Fig **4C**).

A major inference from the two-phase model is that P-E connectivity in baseline conditions primes differential gene expression upon global connectivity loss, highlighting its predictive power for cancer gene dysregulation. To test this hypothesis, we developed an “initial” machine learning model using connectivity and other omics landscapes in mucosa to predict gene expression changes in advanced stages (Fig **4D**). The predicted fold changes exhibited moderate but highly significant correlation for the test gene set in both polyps (r=0.50) and adenocarcinoma (r=0.44) (Fig **4E**, **S6B**). The “initial” model performed comparably to a “differential” model, which was trained using the fold change of the epigenetic landscapes during malignancy progression, suggesting that the baseline levels and alterations of the epigenetic landscapes explained a comparable portion of cancer gene dysregulation (Fig **4F**, **S6C**).

Interestingly, when the baseline of the ‘initial’ model was changed from mucosa to polyp, which molecularly more closely resembles adenocarcinoma, the prediction accuracy significantly worsened (Fig **S6D-F**). This suggests that the predictive information in the epigenetic landscape diminishes as oncogenic progression advances. We identified P-E connectivity as the most critical predictive feature in the ‘initial’ model by sequentially removing each feature, reinforcing its pivotal role in the two-phase model (Fig **S6G**). Analysis of feature importance revealed distinct predictive factors by the “initial” and “differential” models. While the “initial” model suggested that up-regulation was predicted by high P-E connectivity and low enhancer accessibility in unaffected mucosa, the “differential” model suggested that it was associated with their loss and gain, respectively, in advanced stages (Fig **4G**, **S6H, I**).

Recent studies on cancer dysregulation have inferred that oncogenic mutations converge on dysregulation of key transcriptional regulators, such as MYC^45^ and E2F^46^, which drive cell proliferation and survival. To test if the two-phase model is a potential addiction mechanism adopted by cancers to gain proliferative advantages, we applied the “initial” model to predict gene dysregulation in other cancer types from the TCGA database^47,48^. We found that the model had generic predictability for significantly dysregulated genes (10/28, AUC > 0.6, Fig **S7**) and the directionality of their differential expression (18/28, accuracy > 60%, Fig **4H**). The prediction accuracy further increased towards nearly 100% for genes showing consistent up- or down-regulation among different cancer types (Fig **4I**). The genes with accurately predicted up-regulation were highly enriched in cell cycle and DNA maintenance pathways (Fig **4J**). These results suggest that a significant portion of conserved cancer gene dysregulation may be explained by the transition of P-E connectivity to the activation phase, including the up-regulation of key transcriptional addiction pathways, such as cell proliferation.

### Two-phase model predicts gene- and cancer-specific sensitivity to epigenetic interventions

In the two-phase model, genes with high P-E connectivity are associated with high stability and thus more transcriptionally resilient to connectivity loss. This model predicts specific transcriptional outcomes driven by connectivity interventions: first, the genome-wide perturbation of the P-E connectome will result in gene-specific expression changes, where low-connectivity genes will be selectively prone to down-regulation; second, high-connectivity genes will be sensitized to perturbations in polyps and cancer due to their shift towards the activation phase along with global connectivity losses (Fig **5A**).

**Figure 5.**
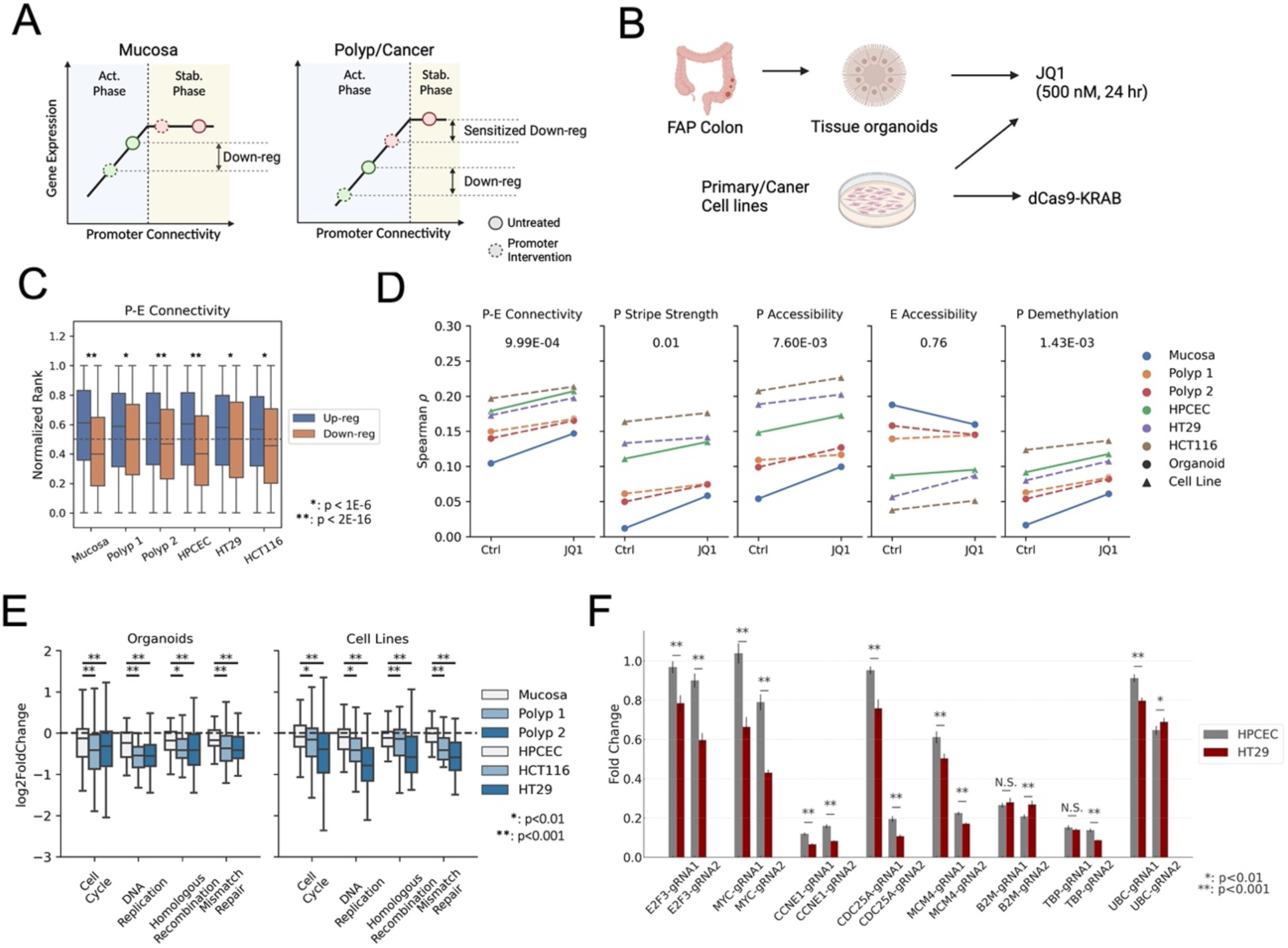
Two-phase model predicts gene- and polyp/cancer-specific sensitivity to inhibitions. (A) Schematic representing the prediction model: Low connectivity genes (green dot) are vulnerable to perturbations in the activation phase, independent of overall connectivity levels. Conversely, genes with high connectivity exhibit stage-specific sensitivity to perturbations as they approach the phase transition threshold due to connectivity loss. (B) Diagram of experimental designs for assessing gene expression sensitivity to various interventions. (C) Distribution of promoter-enhancer (P-E) connectivity levels in unaffected mucosa for genes upregulated or downregulated after JQ1 treatment. p-values for significance from the Mann–Whitney U test are indicated. (D) Spearman correlation between structural and epigenetic features in mucosa and gene expression levels in cell lines and organoids pre- and post-JQ1 treatment. p-values for significance from the Wilcoxon signed-rank test are shown. (E) Expression fold change distributions for genes in specified pathways following JQ1 treatment. p-values for sample differential responses from the Wilcoxon signed-rank test are denoted. Error bars represent confidence intervals. (F) Differential gene expression following Cas9-KRAB-mediated repression via two guide RNAs in primary human colon epithelial cells (HPCEC) and the HT29 colorectal adenocarcinoma cell line. The significance of differential responses was assessed using a two-sample t-test (N=8). Error bars represent the standard error.

To validate these predictions, we first examined the transcriptomic response of colon samples to JQ1 treatment (Fig **5B**). JQ1 is a specific and potent inhibitor of BET family members^49–52^, such as BRD4, which plays a crucial role in recruiting the mediator complex that bridges promoter-enhancer interactions^53,54^. In primary and colon cancer cell lines, as well as in organoids derived from mucosa and polyp tissues, genes up- or down-regulated by JQ1 treatment were consistently associated with high and low initial P-E connectivity, respectively (Fig **5C**). Furthermore, correlations of transcription levels with P-E connectivity and other indicators of promoter activity, but not with the enhancer context, were increased with JQ1 treatment (Fig **5D**, **S8A**). These results resembled the events during malignancy progression (Fig **4A, B**), suggesting that gene expression alterations induced by JQ1 could be explained by the two-phase model.

By comparing the P-E connectivity distribution of down-regulated genes in normal versus polyp/cancer samples, we found a significant increase in the fraction of high-connectivity genes in disease samples (Fig **S8B**). This result matched the malignancy-specific sensitivity of highly connected genes to perturbations predicted by the two-phase model. Interestingly, genes that were selectively down-regulated in polyps and cancer cells were enriched in cell cycle and DNA damage repair pathways (Fig **S8C**). These pathways responded to JQ1 with a significantly higher fold decrease in disease samples (Fig **5E**), suggesting that while cell proliferation genes are commonly up-regulated during oncogenesis (Fig **4J**), their susceptibility to transcriptional perturbations also increases with impaired P-E connectivity.

To test whether the cancer-specific perturbation sensitivity can be targeted with a gene-specific strategy, we applied CRISPRi^55^ to target the promoters of five proliferation genes (E2F3a, MYC, CCNE1, MCM4, CDC25A), which were highly connected and correctly predicted to be upregulated in both polyps and adenocarcinoma (Fig **S5A**, Table **S1**). Comparing transcriptional responses in primary colon epithelial cells (HPCEC) and colon cancer cells (HT29), we found that each of the genes were consistently repressed with a larger effect size in the cancer cell line (Fig **5F**). In contrast, for the three reference genes (B2M, TBP, UBC), neither or only one of the two guide RNAs targeting each gene showed increased repression fold changes in cancer. These results were reproduced by delivering dCas9-gRNA ribonucleoprotein complexes via electroporation, replacing the lentiviral-delivered Cas9-KRAB cassette (Fig **S9A**). By contrast, gene repression and deletion using the exon-targeting Cas9/dCas9 resulted in similar degrees of down-regulation between normal and cancer cell lines (Fig **S9B, C**), suggesting that the observed difference in repression efficiency was specific to the promoters and not confounded by the delivery efficiency of the system between cell lines. Taken together, consistent with the two-phase model prediction, proliferation genes in cancer showed increased susceptibility to promoter inhibition.

## Discussion

In this study, we have provided a comprehensive and integrative analysis of P-E connectivity in conjunction with the transcriptional and epigenetic state of regulatory elements during the early stages of CRC development. Our high-resolution chromatin conformation data, facilitated by multi-restriction digestion, revealed a large number (>250,000) of dot interactions and architectural stripes associated with active regulatory elements, such as promoters and enhancers. This represents a pivotal departure from previous descriptions of chromatin architecture that primarily focused on CTCF loop structures and large domain regions^30,56–58^. Our findings indicate that most P-E loops co-existed with stripe formation, suggesting that these interactions are highly dynamic, occurring as either anchor sliding over the intervening chromatin. This challenges the traditional view of stable P-E loops and aligns with the recent proposal of the “hub” model^59,60^, which describes the close vicinity but not tight looping of cis-regulatory hubs. Importantly, our study shows that P-E connectivity has a significant inference to gene expression dysregulation in cancer, underscoring its fundamental role in gene regulation.

Our results elucidate the distinct roles of P-E connectivity and enhancer activity in CRC progression. During the transformation of polyps and adenocarcinomas, we observed that up-regulation of gene expression was often correlated with increased activity of the neighboring enhancer contexts (Fig **4A,G**). Conversely, P-E connectivity diminished for most genes, irrespective of their expression changes (Fig **S5C**). This intriguing dichotomy underscores the existence of specific regulatory mechanisms governing connectivity that are integral to gene expression and CRC development. The attenuation of P-E connectivity is primarily linked to the reduction in stripe activity on gene promoters (Fig **3C**). However, the precise mechanisms driving the disproportionately high loss rate of connectivity of promoters among CREs remain elusive (Fig **S4B**). Recent investigations have spotlighted the role of chromatin binding-factors like Pol II, the mediator complex, and YY1 in sustaining P-E interactions^53,61,62^. Additionally, it has been posited that the cohesin complex retention on promoters is modulated by general transcription activities^2,63^. This suggests that a myriad of transcriptional regulators could influence P-E connectivity, thus potentially leading to new targets for therapeutics.

The global loss of connectivity at promoter regions in advanced stages suggests altered dynamics of TFs, associated with a weakened ability to maintain proximity with distal regulatory elements. This altered dynamic is evident in the elevated sensitivity of cell proliferation gene promoters to perturbations in cancer cells, as demonstrated by our inhibition assays. By developing strategies that specifically target these vulnerable promoters, it may become possible to disrupt key oncogenes in a disease-specific fashion, thereby offering a novel and potentially effective approach to early CRC interventions.

We observed an intricate and non-linear relationship between P-E connectivity and transcription. The similar degree of connectivity loss was associated with both up- and down-regulation, and our analysis revealed that this gene-to-gene variation was correlated with their connectivity levels established in the unaffected mucosa prior to cancer development. We propose that this non-linear relationship is a result of a transition in the connectome-transcriptome relationship during CRC development. In the unaffected FAP mucosa, transcription levels are not rate-limited by high P-E connectivity. This redundancy is consistent with previous observations of low correlation between P-E interaction and gene expression^12,13,22,23^. However, our results indicate that the scenario is altered in polyps and cancers, where the loss of connectivity becomes a limiting factor for transcriptional regulation and thus correlates with gene expression. Thus, the cis-regulatory connectivity plays a pivotal role in gene dysregulation associated with cancer progression.

Based on the two-phase model, we reasoned that genes with high and low P-E connectivity at baseline condition would be primed for up- and down-regulation, respectively, upon global connectivity loss or perturbation. This insight was corroborated by the correlation between transcriptional alterations caused by BRD4 inhibition through JQ1 treatment and the initial P-E connectivity levels in unaffected mucosa. Notably, while previous research has reported a comparable number of up- and down-regulated genes in response to JQ1 treatment, the mechanistic underpinnings of widespread gene up-regulation following the inhibition of BRD4, a general transcriptional activator, remained elusive^64,65^. Our study provides a possible explanation, suggesting that the up-regulation of genes can be attributed to their promoters’ tolerance to BRD4 inhibition compared to the rest of the genome.

Notably, we identified early established high P-E connectivity in unaffected mucosa tissue as a hallmark of gene up-regulation during oncogenic progression. Our pan-cancer analysis suggests that this hallmark is significant in multiple cancer types, and this finding was particularly pronounced in key transcriptional regulators of proliferation, such as E2F and MYC (Fig **S6A**). Previous studies have often described the upregulation of MYC and E2F as an outcome of genetic mutations or alterations in their upstream regulators^45,66–68^. However, our results suggest an alternative perspective, where the global remodeling of regulatory connectivity plays a significant role in their frequent up-regulation in cancers.

Interestingly, our findings suggesting that P-E connectome remodeling serves as a positive driver in oncogenesis are in contrast with a recent topological study of colon cancer, which proposed a tumor-suppressive effect of large-scale architectural reorganization^56^. This apparent discrepancy likely reflects the distinct influences of macroscopic chromatin structures in previous studies and the microscopic P-E interactions in cancer progression identified in our study. While compartmental remodeling within repressive domains coincided with their hypomethylation and gene repression, we observed a concurrent loss of fine-scale connectivity among active regulatory elements, shifting the global transcriptional balance.

One limitation of our study is its reliance on colorectal tissues from a relatively small cohort of FAP patients. Previous studies have shown that even seemingly unaffected intestinal mucosa in FAP patients displays deregulated proliferation compared to tissues from genetically normal individuals^69–71^. Whether this predisposition towards tumorigenic transformation is associated with chromosomal conformational remodeling similar to the changes we observed in subsequent stages of progression remains to be explored. Additionally, while our two-phase model was robust across several cancer types, it did not effectively predict gene dysregulation for certain cancers such as kidney carcinoma and myeloid leukemia (Fig **4H**). This discrepancy may indicate cell-type-specific variations in P-E connectivity, underscoring the need for comparative studies involving these cancers and their respective healthy controls. Future research should expand to more diverse cohorts and cancer origins to fully assess the complex relationship between regulatory connectivity and gene dysregulation proposed by our two-phase model.

In summary, our study offers valuable insights into the complex interplay between 3D genome architecture and gene regulation during the early stages of CRC progression. We provide a unique resource of fine-gauge regulatory architecture that has not been extensively explored in previous cancer chromatin conformation mapping studies. By comprehensively tracing the dynamic changes in P-E connectivity and their impact on gene expression during early CRC development, we have identified potential new paths for therapeutic interventions. For example, by restoring normal P-E connectivity, it may be possible to interfere with the gene dysregulation events during CRC progression. Further dissection of mechanisms underlying altered cis-regulatory connectivity during the disease development could identify transcriptional regulators that trigger cancer-specific suppressions of oncogenes, opening up new avenues for treatment.

## Author Contributions

Conceptualization, Y.Z., M.P.S.; mHi-C and RNA seq, Y.Z.; ATAC seq: K.P., A.K.W.; EM seq, H.L.,G.K., A.J., D.L., Z.S.; Data Analysis, Y.Z.; Sample Collection, R.L., M.M., A.H.,U.L., E.D.E., R.C., J.S.; Organoid Resources: S.W.; Original Draft, Y.Z., M.P.S., Review and Editing, All Authors; Funding Acquisition, A.K., C.C., E.D.E, J.M.F., W.J.G., M.P.S.; Supervision, E.D.E., A.K., J.M.F., C.C., W.J.G., M.P.S..

## Acknowledgements

This study is supported by NCI funding 1U2CCA233311-01. Illumina sequencing of mHi-C, ATAC seq, and RNA seq were performed by the Stanford Genomics center. The data was generated with instrumentation purchased with NIH funds S10OD025212 and 1S10OD021763. Some illustration figures were created with BioRender.

## Ethics Declarations

G.K., A.J., D.L. and Z.S. are employees and shareholders of Ultima Genomics. M.P.S is cofounder and scientific advisor of Personalis, Qbio, SensOmics, January AI, Mirvie, Protos, NiMo, Onza and is on the advisory board of Genapsys. Aziz Khan has affiliations with Biogen (consultant), SerImmune (SAB), RavelBio (scientific co-founder and SAB) and PatchBio (SAB). E.E. is an employee and stockholder of Invitae, advisor and stockholder of Taproot Health and Exir Bio. William Greenleaf has affiliations with Guardant Health (Consultant/SAB), Protillion Biosciences (Scientifica Co-Founder), and 10x has licenced patents associated with ATAC-seq.

## Data Availability

Raw and processed sequencing data are available at NCBI Gene Expression Omnibus (GEO), accession number GSE207954, dbGAP (phs002371), and on the NCI Human Tumor Atlas Network portal (RRID: SCR_023364).

## Methods

### Sample Collection

FAP tissues were collected at the time of partial or full colectomies from patients. Immediately following colectomy, patient-matched non-neoplastic colorectal mucosa, adenomatous polyps, and adenocarcinomas were snap-frozen and preserved in liquid nitrogen. One FAP adenocarcinoma (A001-C-007) was embedded in an optimal cutting temperature compound (OCT) before being stored at −80°C. Six sporadic CRCs were obtained from the Stanford Tissue Bank. Tissues were examined for histopathology to confirm their disease states. All collection procedures were conducted under IRB protocol 47044.

### Organoid Culture

Tissue samples were collected from patients and processed for organoid generation according to the protocol detailed in Pleguezuelos-Manzano et al.^72^. Briefly, samples were collected on ice in a collection medium (advanced DMEM/F12 supplemented with 10 mM Hepes, 1× Glutamax, and 1% penicillin/streptomycin). Tissue samples were washed in collection medium, minced, and digested for 30 minutes at 37°C in 5 mg/mL collagenase type II (SigmaAldrich). Samples were then filtered using a 100 μm strainer, washed five times in collection medium, and plated in Geltrex (ThermoFisher).

Organoids were cultured in a complete medium (advanced DMEM/F12 supplemented with 10 mM Hepes, 1× Glutamax, 1% penicillin/streptomycin, 1x B27 without vitamin A, 10 mM nicotinamide, 1.25 mM N-acetylcysteine, 500 nM A83-01 (Tocris), 10 μM SB202190 (SigmaAldrich), 100 ng/μl Noggin (R&D Systems), 1 μg/mL human recombinant R-spondin (Stemcell), 0.3 nM Wnt-FC (Immunoprecise), 50 ng/ml EGF (Shenandoah Biotechnology, Inc), 2.5 μM CHIR 99021 (Tocris) and 100 μg/ml Normocin [InvivoGen]). Ten micromolar of Y-27632 was added to the medium for the first three days after seeding.

For drug experiments, organoids were trypsinized and plated at 30,000 cells/well in 24-well plates. After 5-7 days, organoids were incubated in a complete medium containing 500 nM JQ1 for 24 hours. Organoids were then harvested in Cell Recovery Solution (Corning) on ice for 1 hour, washed with PBS, and centrifuged to retrieve cell pellets. Cell pellets were then processed for RNA extraction.

### Cell Lines

HT-29 (ATCC HTB-38) and HCT-116 (ATCC CCL-247) human colorectal cancer cell lines were obtained from ATCC. Cells were maintained in DMEM-F12 (ThermoFisher 11320033) with 10% FBS and 1% penicillin/streptomycin. Primary human colonic epithelial cells (Cell Biologics H-6047) were maintained in the Epithelial Cell Growth Medium (Cell Biologics H6621) as suggested by the vendor. All cell experiments were performed before reaching 10 population doublings.

For drug experiments, cells were seeded in 6-well plates with 50% confluence. After one day, cells were grown in a complete medium containing 500 nM JQ1 for 24 hours. Cell pellets were then trypsinized for collection and processed for RNA extraction.

### CRISPR/CRISPRi Assays

For stable expression of dCas9-KRAB, pHR-SFFV-KRAB-dCas9-P2A-mCherry (Addgene 60954) was transfected into HEK293T cells using the Lenti-X Packaging Single Shots (Takara Bio 631275), following the manufacturer’s protocol. Assembled viral particles were harvested after 72 hours of incubation and collected by filtering the culture media through a 0.45 µM filter. Viral titers were determined using the Lenti-X GoStix Plus (Takara Bio 631280). Viral infections were conducted with a MOI (multiplicity of infection) of 10, where 2.0E5 to 1.0E6 cells were incubated in full culture media containing the viral particles and 8 µg/mL polybrene for 48 hours. Positive cells were selected based on high expression of mCherry by FACS sorting (FACSAria II, BD Bioscience). Cells with stable cassette expression were selected by a second sorting performed 2-3 weeks after the initial.

For delivery of gRNA or Cas9/dCas9-gRNA complex, synthesized crRNA:tracrRNA duplex (IDT) were transfected into 1.0E5-2.0E5 cells using the 4D nucleofector X unit (Lonza), with or without pre-incubation with equal molar Cas9/dCas9 protein (IDT 1081058,1081066). We followed the protocol provided by IDT for the Lonza Nucleofector System. Nucleofection of the cells used the following kits and programs: HPCEC - P3/CM137; HT29 - SF/FF137. After nucleofection, cells were seeded in 96-well plates and collected for downstream analysis after 48 hours. To obtain statistical robustness, each experiment was repeated with 2-4 trials with two replicates in each trial.

### Multi-digested Hi-C (mHi-C)

Multi-digested Hi-C (mHi-C) was performed as a derivative of Tri-HiC, a high resolution modified Hi-C protocol ^18,31^, with minor modifications. Initially, 5-10 mg of snap-frozen tissue was placed into a tissueTUBE-TT05 (Covaris 520071) and cryopulverized using the Covaris CP02 cryoPREP Automated Dry Pulverizer, following the manufacturer’s procedure. The pulverized tissue was then subjected to freeze substitution^73^ by submerging it in 1 ml of −80°C 0.01% formaldehyde (ThermoFisher 28906), 97% ethanol, and 2% water. Following this, samples were incubated on dry ice for 3 hours at a rotor spinning speed of approximately 100 rpm. They were then placed in a CoolCell Container (Corning) and transferred to a −20°C freezer for overnight incubation. On day 2, the container was moved to a 4°C cold room and spun on a rotor at approximately 100 rpm for 1 hour to bring the sample temperature above the freezing point.

Subsequently, the tissue samples were separated from the ethanol solution by centrifuging at 300 g for 5 minutes in a 4°C microcentrifuge. Crosslinking was carried out by incubating the samples with 1 ml of 1% TBS-formaldehyde for 10 minutes at room temperature. The solution was then quenched by adding 80 μl of 2.5 M glycine and incubated for an additional 5 minutes. The samples were centrifuged, washed once with 1 ml of TBS (pH 7.5), and resuspended in 250 μl of Hi-C lysis buffer (10 mM Tris-HCl, pH 8.0, 10 mM NaCl, 0.2% Igepal CA630) with an additional 50 μl of proteinase inhibitor cocktail (Sigma P8340). Nuclei extraction was performed on ice by squeezing the samples 15-20 times with 1.5 ml disposable pellet pestles (Fisher Scientific 12-141-368).

The crude suspension was then centrifuged at 1500 g for 5 minutes at 4°C, resuspended in 800 μl of Hi-C lysis buffer, and passed through a 100 μm strainer (Sysmex). After another centrifugation, the purified nuclei were resuspended in 170 μl of 10 mM Tris-HCl containing 0.5% Triton X-100 (Sigma 93443). This was followed by incubation at room temperature with rotation for 15 minutes. Ten microliters of 1% SDS, 20 μl of Cutsmart buffer (NEB), 3 μl each of HinP1I (NEB R0124S), DdeI (NEB R0175L), CviAII (NEB R0640L), FspBI (ThermoFisher ER1762), and 0.6 μl of MseI (NEB R0525M) were added to the suspension in the indicated order. The mixture was then incubated at 25°C and 37°C for 2 hours each, with rotation.

To halt the restriction digestion, the suspension was incubated in a 62°C heating block for 20 minutes, followed by cooling down. End repair was carried out by adding 30 μl of a solution containing 0.5 mM biotin-14-dATP (Active Motif 14138), 0.5 mM biotin-14-dCTP (AAT bio 17019), 0.5 mM dTTP, 0.5 mM dGTP, and 4 μl of Klenow DNA polymerase (NEB M0210L) to the mixture. This was then incubated for 1 hour at 37°C with rotation. For ligation, a 750 μl solution containing 1x NEB T4 DNA ligase buffer (NEB B0202), 120 μg of BSA (ThermoFisher AM2616), and 2000 U of T4 DNA ligase (NEB M0202M) was added. The mixture was incubated at room temperature for 90 minutes, followed by 4°C overnight, and then room temperature for an additional 60 minutes with rotation.

Reverse crosslinking was performed by centrifuging the mixture at 1500 g for 5 minutes. The supernatant was then replaced with a mixture of 300 μl of 1x T4 ligase buffer, 30 μl of 20 mg/ml proteinase K (ThermoFisher 25530049), 30 μl of 10% SDS, and 40 μl of 5 M NaCl. This suspension was then incubated at 66°C for 4 hours. The DNA content was purified by phenol-chloroform extraction and resuspended in 20 μl of 10 mM Tris-HCl.

To generate the mHi-C sequencing library, 300 ng of purified DNA was tagmented with 2.5 μl of Tn5 transposase (APExBIO K1155, discontinued) loaded with equimolar Mosaic Ends containing Illumina Nextera i5 and i7 extensions, according to the manufacturer’s protocol. The tagmentation was performed in a 100 μl buffer containing 10% DMF, 10 mM Tris-HCl, and 150 mM NaCl, at 55°C for 10 minutes. The product was then column purified (Zymo D4014) and PCR amplified for 2 cycles using the NEBNext master mix (NEB M0544L) with Illumina Nextera primers and conditions. Biotin enrichment was then performed by adding 20 μl of Dynabeads MyOne Streptavidin C1 (ThermoFisher 65001) and incubating at room temperature for 30 minutes with rotation. The magnetic beads were washed three times with 1x wash buffer (10 mM Tris-HCl pH 7.5, 1 mM NaCl, 0.5 mM EDTA) and once with 10 mM Tris-HCl. Final libraries were obtained by amplifying the beads with an additional 8 cycles of PCR, followed by purification with SPRI (Beckman B23318) size selection at a 0.5x-1.1x range. The 33 samples were combined into 2 pools and sequenced using 2 NovaSeq (Illumina) S4 200 cycle flow cells.

### Real-time PCR/ RNA seq

Total RNA was extracted from approximately 5-10 mg of frozen tissues or approximately 1.0×1051.0×105 to 1.0×1061.0×106 cells from organoid or cell culture, using the Zymo Quick-RNA Miniprep (Zymo R1054), according to the manufacturer’s instructions. After purification, DNA digestion was carried out using the DNA-free DNA Removal Kit (ThermoFisher AM1906).

For cDNA synthesis, up to 1 μg of total RNA was processed using the Superscript IV reverse transcription system (ThermoFisher 18091050), with Oligo dT provided in the kit serving as the primer. For RT-PCR, 50 ng of synthesized cDNA was mixed with 10 μL of TaqMan Fast Advanced Master Mix (ThermoFisher 4444557) and 1x primers, and then examined in the QuantStudio 6 Flex system (ThermoFisher). Relative gene expressions were normalized against the internal expression of GAPDH, utilizing the double delta CT method.

Sequencing libraries of mRNA were prepared from 200 ng to 1 μg of total RNA using the NEBNext Ultra un-stranded preparation kit (E7775S, E7490S), in accordance with the manufacturer’s protocol. Samples were sequenced on a NovaSeq S1 flow cell for 50 bp pair-end sequencing, resulting in an average of 86.3 million raw paired reads per sample.

### ATAC seq

ATAC seq was conducted following the latest ENCODE tissue protocol, as described^61^. Sequencing was performed on a NovaSeq S1 flow cell using 50 bp pair-end sequencing, resulting in an average of 53.2 million unique fragments mapped for each sample.

### EM seq

Enzymatic Methyl seq was executed as described^74^. Libraries were constructed using the NEBNext Enzymatic Methyl-seq Kit (NEB), following the manufacturer’s guidance. Sequencing was carried out using the novel ultrahigh throughput UG-100 (Ultima Genomics) sequencer.

### mHi-C data processing

Initial processing of mHi-C data was executed utilizing the distiller pipeline (https://github.com/open2c/distiller-nf) with default parameters configured for the SLURM cluster. Deduplicated pair files were then input into Juicer pre^37^ to generate KR-balanced .hic matrices at resolutions of 200, 500, 1k, 2k, 5k, 10k, 20k, 50k, 100k, 250k, 500k, and 1mbp, employing a quality score filter of 30. In order to generate piled-up master matrices for various stages and all samples, pair files were initially merged and sorted utilizing pairtools (https://github.com/open2c/pairtools).

Stripe calling was carried out as previously described^18^, with minor modifications to the parameters. We noted an enrichment of mappable reads at open chromatin regions, raising the possibility of false stripe signal detection by measuring raw read count over-representation. To address this, the stripe calling algorithm normalized long-range contacts against read mappability at each locus, evaluated by distal interactivity-independent self-ligation events. Specifically, long-range (>1.5 kb) and short-range (<1 kb, +/− orientation) mapped read pairs were separated into two .bam files using awk and samtools. Bedtools was then utilized to map these reads to two binning bed tracks; a local one with a 2 kb window and a background one with a 50 kb window, both featuring a 100 bp sliding size. Assuming the over-representation sourced from mappability was proportional between long- and short-range contracts, the expected count number for each bin was calculated as (long_bg_ / short_bg_) × short_local_. Using MACS2 bdgcmp -m qpois, the local long-range read count for each bin was examined for statistical significance of enrichment against the expected number. The log fold change signal (stripe strength) was then calculated with the same formula by inputting the actual and expected values into MACS2 bdgcmp -m logFE. To prevent NaN errors, a pseudo-count of 1 was added.

Post-determination of stripe q values, each 100 bp bin was counted for the number of samples demonstrating significance (FDR < 0.01). Bins with at least three sample hits were deemed significant. These bins were then merged, and only windows with a minimum size of 500 bp were included as final stripe anchors. Stripe peaks overlapping with ENCODE blacklist regions were removed. To mitigate gender variations among patients, only autosomal chromosomes were included for downstream analyses.

Loop calling was performed using the HiCCUPS algorithm from Juicer tools^37^ with the following parameters: -r 500,1000,2000,5000,10000 -f 0.1 -p 4,2,2,2,2 -i 20,10,10,6,6 -t 0.1,1.25,1.75,2 -d 2000,2000,4000,10000,20000. Given that library complexity significantly affects loop calling power, the analysis was not performed for each individual sample. Instead, it was executed on pooled libraries of 1) all samples, 2) all mucosa, 3) all polyps, and 4) all adenocarcinomas. Post-processed loop pixels from all profiles at various resolutions were subsequently merged in the order of high resolution > low resolution from combined> mucosa > polyp > adeca. A loop with lower priority was filtered if both anchors overlapped with a higher priority loop. This master loop list was then applied to each sample to execute individual loop quantification. Loop strengths were calculated by dividing read counts in the identified loops by the expected count from the donut background and log transforming the results. A pseudo-count of 1 was added as necessary to prevent NaN errors. For stage-specific counting, loops with average loop strengths greater than 1.2-fold in samples of the specified stage were regarded as positive.

For annotations for stripe/loop anchors, features were mapped against the Ensembl regulatory build^36^ and TSS from Gencode using Bedtools. If an anchor overlapped with multiple features, the primary annotation adhered to the following order: promoter/TSS (within −1.5 kb to +0.5 kb of any Gencode transcript), active enhancers (defined by H3K27Ac marks in the Ensembl regulatory build), CTCF binding sites, and open chromatin.

For aggregation analysis of loops, APA from Juicer was conducted with the parameters -r 200 -u -n -0 -w 500 -k KR -q 20. The enrichment score was computed as the average intensity of the 10×10 center pixels (2 kb) against the mean of the 100×100 pixels from the bottom left. For the aggregation of stripes, the same function was executed with the parameters -r 200 -u -n -0 -w 250 -k KR -q 20. Given the phenomenal distance-dependent interaction decay at the vicinity of stripes, interaction intensities at specific distances (a.k.a. matrix diagonals) were normalized against the average intensity of the distance. The fold enrichment of the aggregated stripes was then calculated by averaging the normalized values in the center 10 pixels. For visualizations of loop and stripe APA, matrices were log-transformed before being plotted onto heatmaps.

For structural variant calling, contact matrices in .mcool format were generated by using the distiller pipeline, using default parameters as described above. These matrices were analyzed by using the PredictSV function in the EagleC package^41^ using the following parameters: --balance-type ICE --output-format full --prob-cutoff-5k 0.8 --prob-cutoff-10k 0.8 --prob-cutoff-50k 0.9999. Identified SVs were visually confirmed on the Hi-C heatmaps.

### ATAC seq processing

ATAC seq results were processed using the ENCODE-DCC ATAC-seq pipeline (https://github.com/ENCODE-DCC/atac-seq-pipeline) with default settings. To generate the integrated peak list from all samples, a 100 bp binning track was created and mapped with the pseudo-replicated peak regions from each sample using Bedtools. Bins with at least three hits were deemed valid, merged, and intervals with a minimum size of 300 bp were included as final peak sites. Peak fold enrichments of samples were then obtained from pipeline-derived fold change bigwig tracks.

### Analysis of P-E Connectivity

To elucidate the connectivity between promoters and their neighboring regulatory context, P-E pairs were determined for all ATAC-seq peaks within a 200 kb distance from target promoters. Interaction zones were defined as being within −1.5 kb to +0.5 kb near the TSS for promoters, and within 500 bp of ATAC-seq peaks for distal enhancers. Contact frequencies between promoter and each target enhancer were calculated using Juicer Straw^37^ to extract read counts at 1000 bp resolution. These raw contacts were normalized against the read mappability of the promoter, which was defined as the KR-normalized contact frequency, or in the case where balanced matrix was not available, the ratio of short-range self-ligation contact (<1.0 kb, +/− orientation) RPKM at the TSS region versus that for the 50 kb neighborhood. Normalization for coverage between samples was achieved by dividing frequencies of each contact by the long-range contact (with a minimum 1.5 kb distance threshold) densities in the 5-50 kb background donut region, in the unit of contacts per 1×1 kb square. The aggregated normalized contacts of the gene with all distal regulatory elements represented the total P-E connectivity of that gene.

The accessibility of interacting regulatory element peaks, referred to as “enhancer accessibility” in the text, was calculated as the log sum of the fold enrichment signals of all ATAC-seq peaks that were involved in calculating the P-E connectivity. This value was extracted from the processed ATAC fold change .bigwig tracks by using pyBigWig, which represented the overall activity of the regulatory context for each gene, and was used for comparisons with P-E connectivity levels. For promoter accessibility, the top two quantiles of mean fold change of the ATAC-seq signals in the defined TSS region were retrieved using pyBigWig and log-transformed. For downstream analyses, genes showing non-positive values for P-E connectivity, promoter accessibility, or promoter stripe enrichment in all three stages were removed. For the remaining genes, missing or negative values were replaced with zero.

### RNA seq processing

RNA seq results were processed using Tomas Bencomo’s pipeline (https://github.com/tjbencomo/bulk-rnaseq), which utilizes Salmon for quantifying transcript levels and DESeq2 for identifying differential genes (FDR < 0.1, fold change > 1.3). Transcription levels (TPM) of genes were obtained by summing transcript-based TPM from Salmon output (.rf).

### DNA methylation processing

A total of 21,175,510 CpG sites with measurable methylation ratios were identified across all samples. The methylation degree of features, including mHiC hotspots, ATAC peaks, and gene promoters, were calculated by averaging the methylation percentage for all valid CpG sites within the feature. Regions with average methylation < 25% were classified as demethylated, while those > 40% were considered methylated. Regions with average methylation between 25% and 40% were classified as intermediately methylated and excluded from the methyl versus demethyl analyses. For methylation changes between two stages, regions demonstrating > 15% difference with <0.1 FDR were classified as significantly hypo- or hyper-methylated based on the direction of change. For correlation analyses with other features, the degree of demethylation (100% minus methylation percentage, annotated as “demethylation”) was often used to maintain a positive correlation between methylation degree and regulatory activity.

### Mappability analysis

The mappability of mHi-C, in situ Hi-C, and intact micro-C at annotated regulatory regions was visualized by retrieving their read coverage from a .bw file that compiled only short-range self-ligation contacts, defined as those with an interaction distance of less than 1.5 kb and a “+/−” orientation. To compare the distal interaction signal against mappability, long-range interactions, defined as those with an interaction distance greater than 2.0 kb, were extracted and aggregated similarly. The fold enrichment of interaction was then calculated as the ratio of long-range to short-range interactions at 100 bp resolution. Note that this ratio is equivalent to the “raw” stripe strength before normalization against the average enrichment of the local background.

### Principal component analysis

PCA analyses of mHi-C and RNA seq were performed using the Python sklearn.decomposition.PCA package. For P-E connectivity, analysis was conducted using either untransformed or scaled-by-sample matrices.

### Motif enrichment analysis

For each feature (P-E connectivity, enhancer accessibility, TSS accessibility, TSS stripe strength, TSS methylation, gene expression), sequences of promoter regions for genes with the top 10% feature strength in mucosa were extracted using bedtools. Motif enrichment was then calculated using the AME tool in the MEME suite^75^, with the JASPAR 2022 vertebrate motif database serving as the reference. The −log10 p-values for significantly enriched motifs for any of the features were included for hierarchical clustering.

### Gene Ontology

Enrichment analysis of significantly up- or down-regulated genes in pathways was performed and visualized using the WEB-based GEne SeT AnaLysis Toolkit^76^. The method of over-representation was selected to test enrichments in the KEGG pathway against the protein-coding genome. Analysis was performed using default parameters.

### Machine learning

For the “initial” and “differential” models, structural and epigenetic feature values in unaffected mucosa or their fold changes in polyps or adenocarcinoma were compiled for each promoter. For fold change calculation, a pseudo count of 1 was added to connectivity and methylation to avoid zero values. These features were further compiled with the expression levels of the genes in mucosa, the binary binding status of all TFs in the ENCODE ChIP-seq database at their promoters, and the sum of the binary binding status for the TFs in their distal enhancer contexts.

The raw value and their rank-transform were combined, resulting in a total of 1,374 features for each gene. To train the models, 2,800 randomly selected genes were excluded as the test dataset, and the rest were fit to the differential gene expression changes in polyps and adenocarcinoma using a sequential model with three intermediate layers and one dense output. Each layer included 2,048, 512, and 128 neurons in order, and was filtered with a 20% dropout rate. Models were trained for up to 50 epochs, with the final model represented by the iteration that showed the lowest mean squared error for the test dataset. The training and evaluation of the models were performed using the Tensorflow Keras module in Python.

For evaluation of feature importance in the models, the SHAP^77^ package was used for analysis with default parameters. Average feature importance was calculated as the absolute mean of the importance across all genes. Overall directionality was represented by the numerical mean of the importance.

### The Cancer Genome Atlas Program (TCGA) gene expression analysis

A list of differentially expressed genes and their fold changes were obtained from the GEPIA database^47^. The prediction of differential expression was performed using the receiver operating characteristic (ROC) analysis in the Python sklearn package, where predicted differential fold changes from the “initial” model were used as thresholds for the up- and down-regulated genes. For prediction of the directionality, genes were scored by their consistency of dysregulation, averaging their alterations (−1 for down-regulation, 0 for unchanged, and +1 for up-regulation). The correlation between the directionality score and the prediction accuracy was then evaluated.

### Public data usage

An unmasked hg38 genome was utilized as the reference for all analyses. The regulatory build for sigmoid colon (version 20210107) was obtained from Ensembl (http://www.ensembl.org) to facilitate regulatory annotations. Gencode v38 was employed for RNA-seq alignment and defining the positions of transcription start sites (TSS). The ENCODE blacklist^78^ was utilized to exclude problematic regions of the genome from analyses. ENCODE in situ Hi-C (ENCSR123UVP) and intact micro-C (ENCSR477GZK) datasets for the HCT116 cell line were used for comparison with our mHi-C data. Roadmap histone ChIP-seq tracks for colonic mucosa (GSM1112779, GSM916043, GSM916046, GSM916045) and ENCODE CTCF (ENCSR833FWC), Pol II (ENCSR322JEO), RAD21 (ENCSR956UIS), SMC3 (ENCSR149SKU) ChIP-seqs were employed for CRE visualization. Locations of CpG islands were downloaded from the UCSC Genome Browser.

## Supplementary Figures

**Figure S1.**
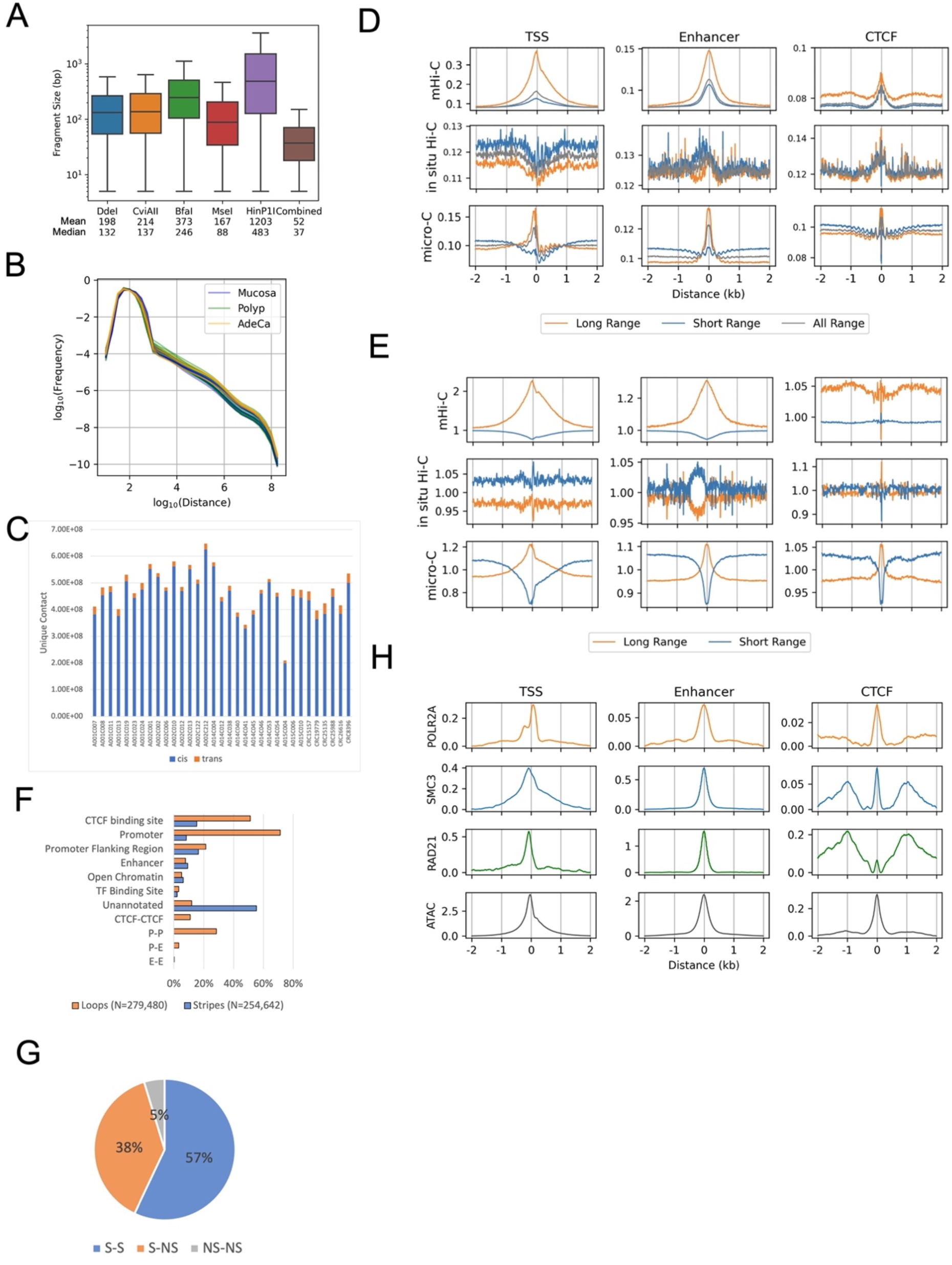
mHi-C delineates interaction features at active CREs. (A) Fragment size distribution of human genome (hg38) digested by the indicated restriction enzymes and their combination. (B) Frequency of intrachromosomal interactions by genomic distance for various sample types. (C) Count of unique interaction contacts identified across samples. (D) Aggregated read intensity of long- (>1.5 kb) and short-range (<1.0 kb) interactions before and (E) after normalizing against total coverage (All range) at distinct CRE categories. (F) Classification and annotation of identified stripes and loops in colon samples by regulatory element types. (G) Proportion of loops composed of two stripe anchors (S-S), between a stripe anchor and a non-stripe anchor (S-NS), and two non-stripe anchors (NS-NS). (H) Aggregated ChIP-seq signals for Pol II, SMC3, Rad21, and ATAC seq fold enrichment at various CRE types in colon samples, sourced from ENCODE data.

**Figure S2.**
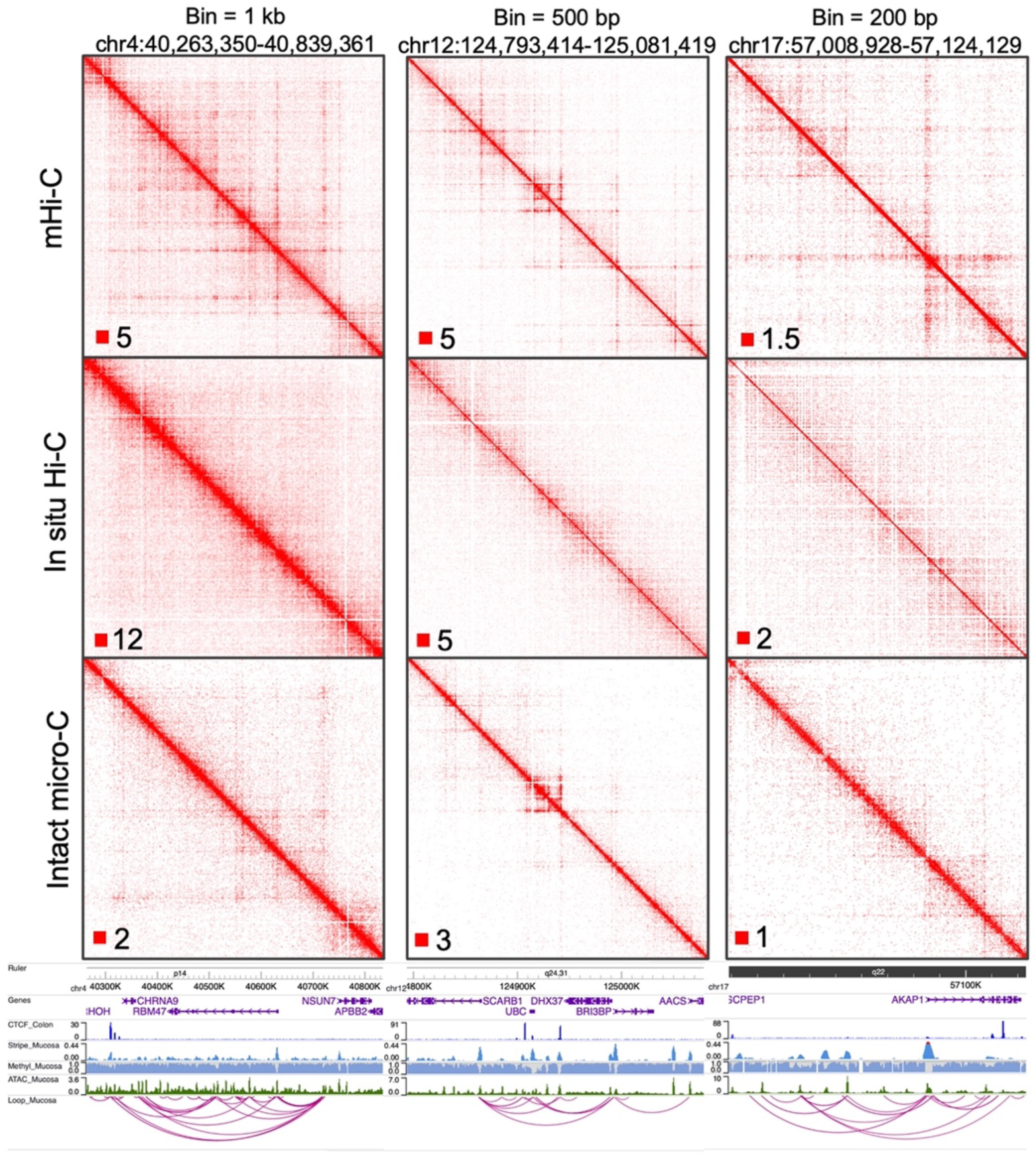
Comparison of mHi-C with in situ Hi-C and intact micro-C at diverse resolutions and genomic loci.

**Figure S3.**
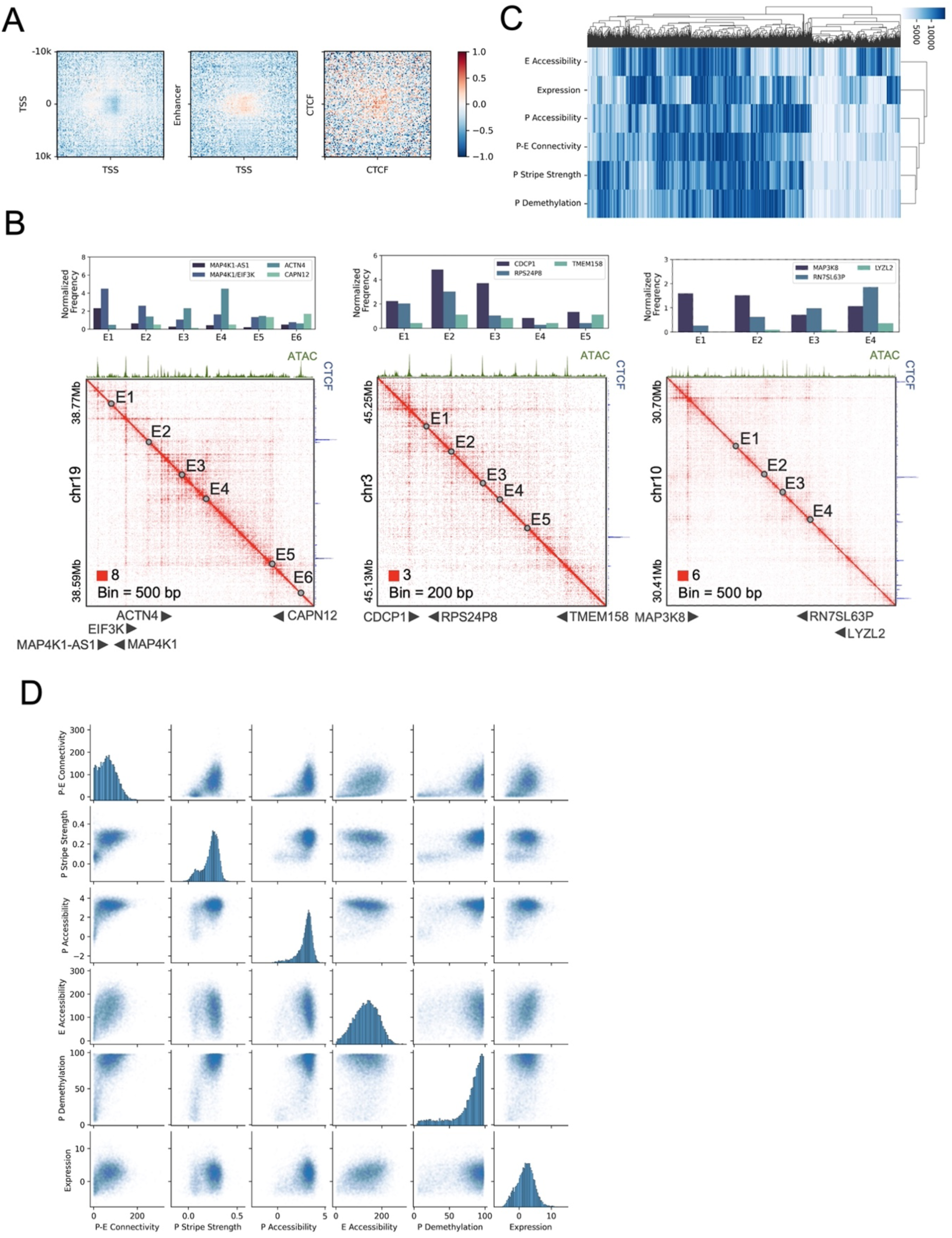
Analysis of the interplay between structural features and epigenetic markers. (A) Heatmaps displaying loop signal intensities as referred to in Fig 2A, adjusted for the effects of stripe strengths at the loop anchors. (B) Contact heatmap at example loci where gene promoters lacking CTCF binding display gene-specific P-E interactions. (C) Hierarchical clustering of genes based on their rankings for various structural and epigenetic features in mucosa samples. Intensity of color corresponds to the strength of the features. (D) Comparative scatter plots illustrating the relationships between different structural and epigenetic features across the examined dataset.

**Figure S4.**
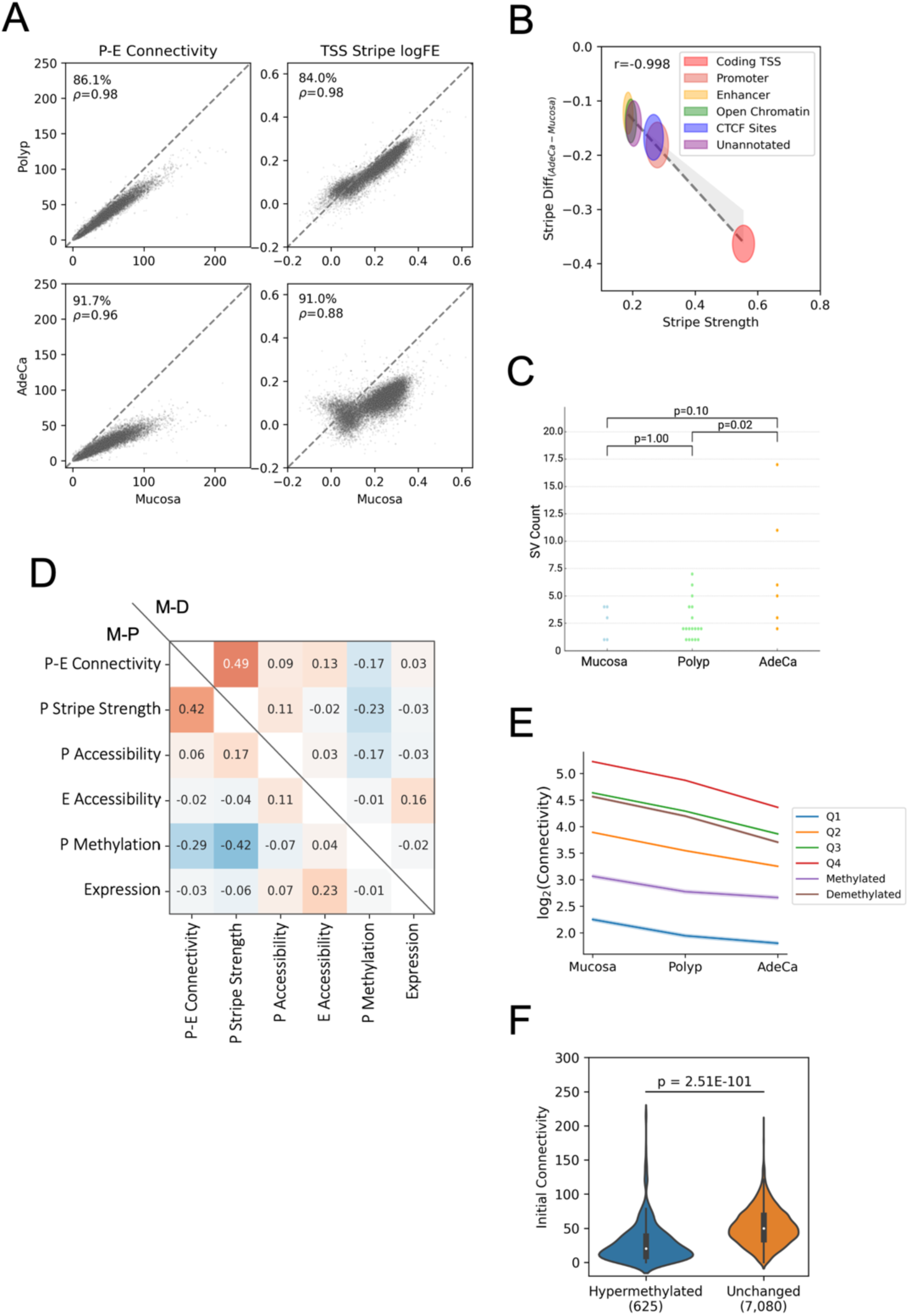
Dynamics of P-E connectivity through colorectal cancer (CRC) progression. (A) Scatter plots depicting the comparative analysis of P-E connectivity and TSS stripe strengths across different CRC stages. The percentage of genes with reduced connectivity (y<x) during progression is indicated for each stage comparison. (B) Correlation between initial stripe strength in mucosa samples and the extent of stripe reduction in adenocarcinoma samples. Each ellipse’s center and radius represent the mean and standard deviation, respectively, for stripes associated with the specified regulatory elements. The dotted line shows the linear regression across the centers of the ellipses. (C) Distribution of SV counts in samples. Significance p values of count differences stages from Mann–Whitney U test are indicated. (D) Spearman correlation matrix detailing the changes (log2 fold change, FC) of structural and epigenetic features between polyps (M-P) and adenocarcinoma (M-D) relative to unaffected mucosa. (E) Average log2 fold change in P-E connectivity for polyps and adenocarcinoma, categorized by promoter methylation status: quantiles (Q1-Q4), demethylated, and methylated, excluding those with minimal hypo- or hyper-methylation. Confidence intervals are depicted as shaded areas behind each line. (F) Distribution of P-E connectivity in mucosa samples for gene promoters that become hypermethylated or remain unchanged in adenocarcinoma. The significance of differences is tested using the Mann-Whitney U test, and the p value is provided.

**Figure S5.**
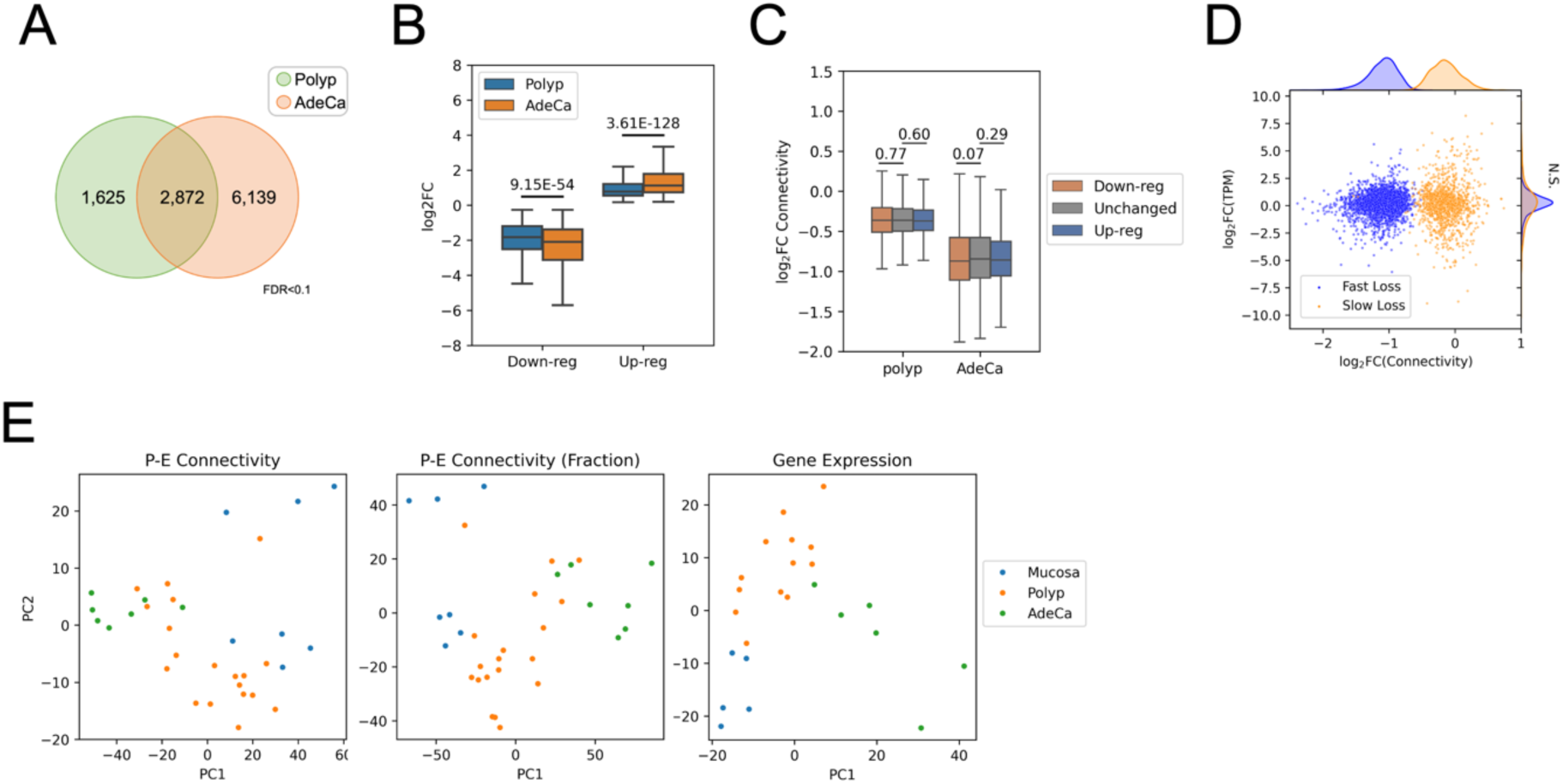
Disconnection between promoter-enhancer (P-E) connectivity and gene expression changes throughout CRC progression. (A) Venn diagram illustrating the commonality of genes with significantly modified expression in both polyps and adenocarcinoma (AdeCa). (B) Log2 fold changes in gene expression for polyps and AdeCa, identifying genes that are consistently up- or down-regulated across both stages. Statistical significance of the difference in fold changes is assessed using the Wilcoxon signed-rank test. (C) Changes in P-E connectivity for polyps and AdeCa relative to mucosa, corresponding to genes with consistent up- or down-regulation. The significance of connectivity changes, as compared to genes without expression alteration, is evaluated using the Mann–Whitney U test. (D) Two-dimensional scatter plots and density distributions correlating the changes in connectivity and gene expression between mucosa and adenocarcinoma. Genes are categorized based on the rate of connectivity loss: fast (blue) and slow (orange), as determined by their feature importance on the first principal component (PC). (E) Principal component analysis (PCA) comparing P-E connectivity, scaled P-E connectivity (normalized against the aggregate sum), and gene expression changes during the stages of CRC development.

**Figure S6.**
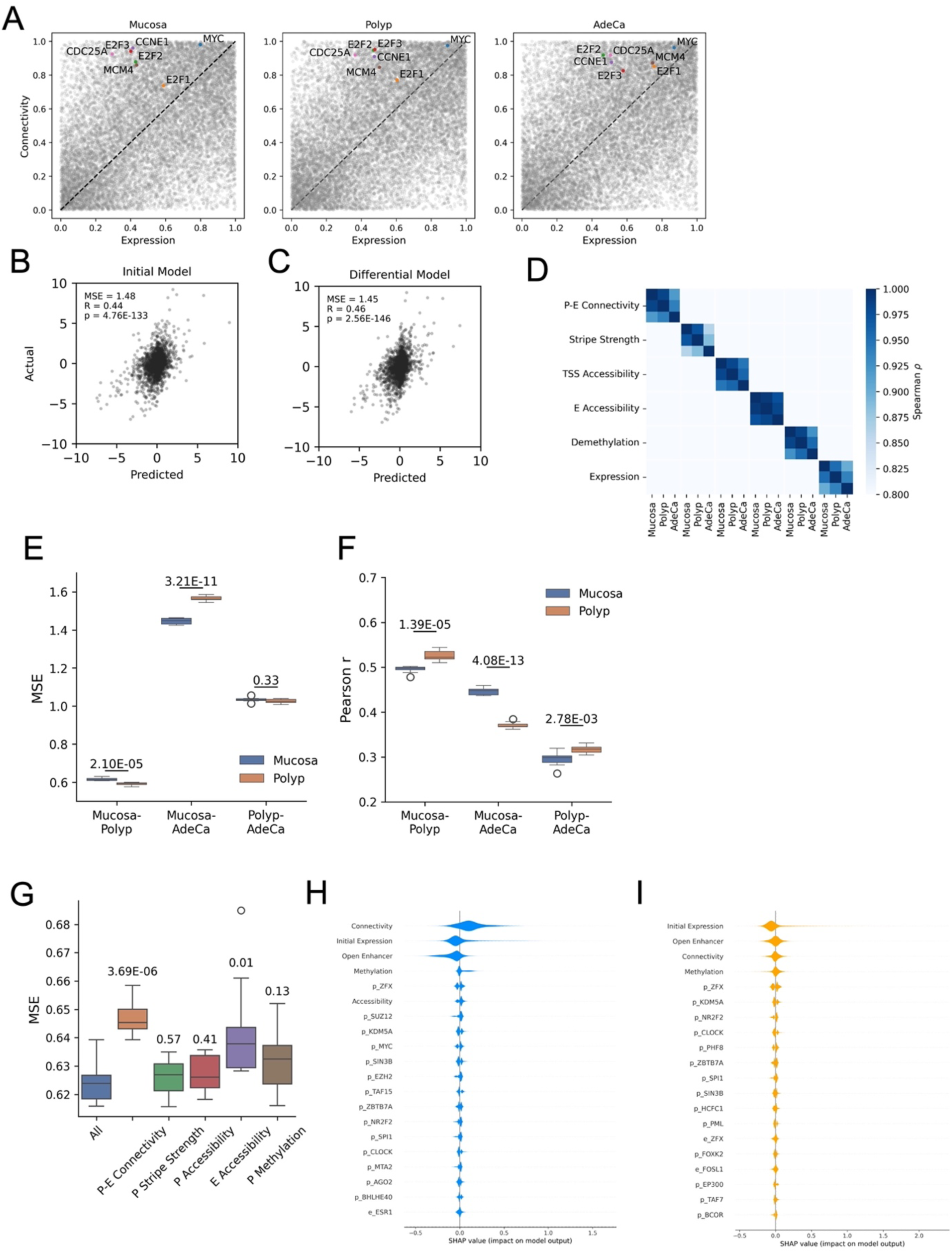
Predictive modeling of gene expression changes based on promoter-enhancer (P-E) connectivity. (A) Distributions of ranks for P-E connectivity and corresponding gene expression levels of key oncogenes and proliferation markers across various cancer progression stages. (B) Model fit assessment for the predicted changes in gene expression in adenocarcinoma using the “initial” model, which utilizes the baseline P-E connectivity. (C) Model fit assessment for the predicted changes in gene expression in adenocarcinoma using the “differential” model, which considers changes in epigenetic landscapes. (D) Spearman correlation matrix showing the similarity of each feature between different stages. (E) Mean squared error (MSE) and (F) Pearson’s r coefficient of the “initial” model for the prediction of gene expression changes in adenocarcinoma compared to the indicated baseline stages. Prediction scores obtained by models trained with mucosa and polyp datasets were compared by using independent t test (N=10). (G) Distributions of minimal mean square error (MSE) of “Initial” model trained with equal or less than 20 epochs (N=10) with the removal of indicated features. Significance p values of differential MSE caused by missing features compared to complete model (All) are evaluated by using independent t-test. (H) The top 20 influential features impacting gene expression predictions in adenocarcinoma, as determined by SHAP (SHapley Additive exPlanations) analysis for the “initial” model. (I) The top 20 influential features for the “differential” polyp model, with features named after transcription factors indicating their binding presence at the promoter (p) or enhancer (e) regions, based on the ENCODE database.

**Figure S7.**
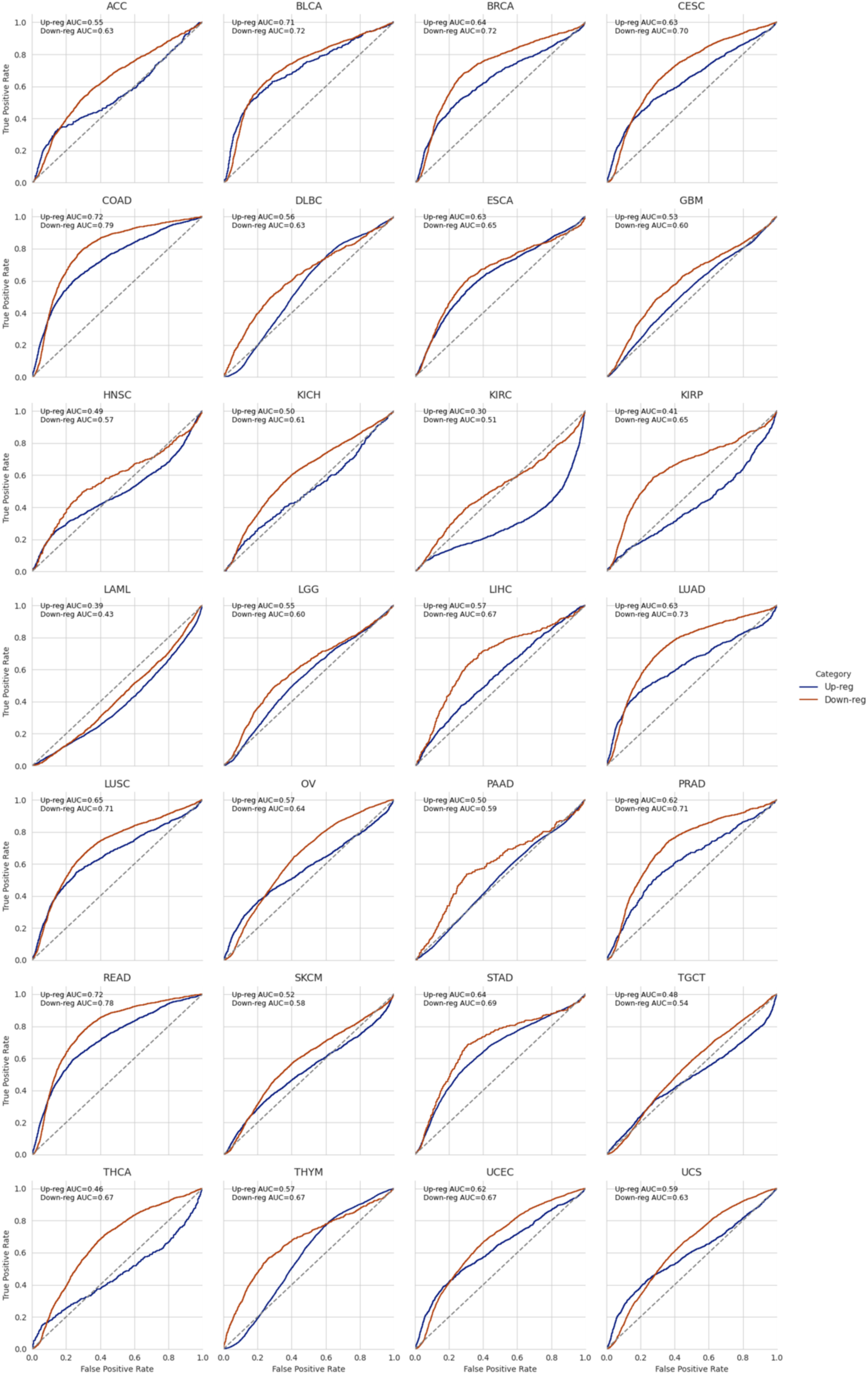
Assessment of predictive accuracy for gene expression changes in various cancer types. ROC curve analysis using gene expression predictions derived from the “initial” polyp model to determine the up- and down-regulation status of genes across different cancer types represented in the TCGA database.

**Figure S8.**
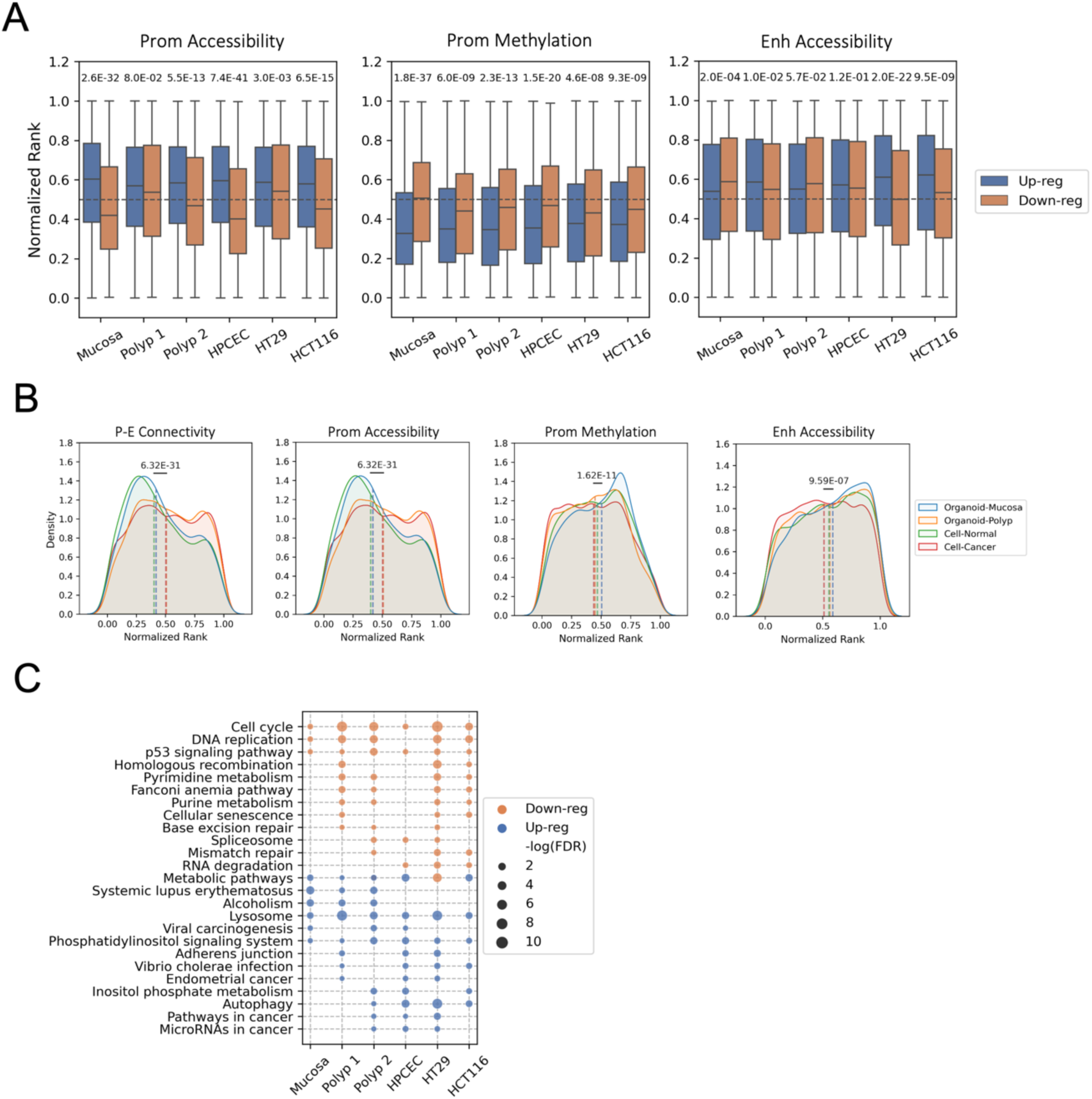
Transcriptomic alterations following JQ1 treatment. (A) Distribution of epigenetic feature ranks in unaffected mucosa for genes that are up- or down-regulated subsequent to JQ1 treatment. P values indicating statistical significance are calculated using the Mann–Whitney U test. (B) Comparative density plots illustrating the differences in feature rank distributions for genes down-regulated in normal tissue (mucosa organoids or primary colon epithelial cells) versus diseased states (polyp organoids or cancer cell lines). P values for statistical significance are derived from the Mann-Whitney U test. (C) Pathway analysis based on ontology for genes that are up- or down-regulated in various samples following JQ1 treatment.

**Figure S9.**
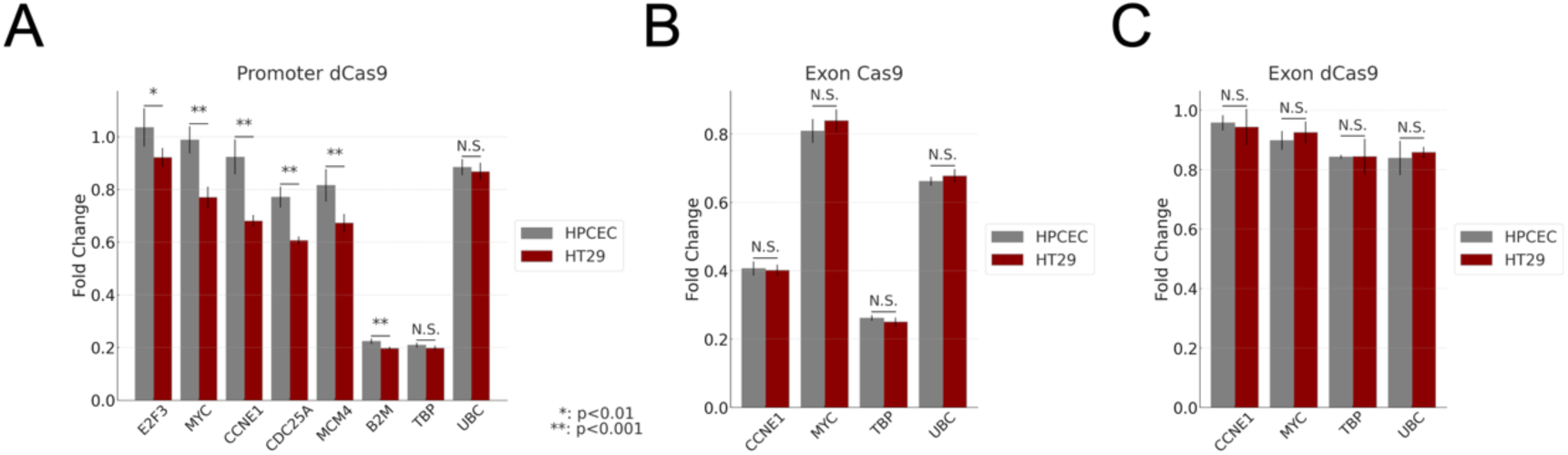
Gene expression changes following Cas9-mediated perturbations in HPCEC and HT29 cells. (A) Changes in gene expression after introducing dCas9-gRNA ribonucleoproteins (RNPs) targeting promoters in wild-type cell lines (N=8). (B) Changes in gene expression after introducing Cas9-gRNA RNPs targeting exons in wild-type cell lines (N=4). (C) Gene expression alterations upon gRNA delivery targeting exons in cell lines stably expressing dCas9-KRAB (N=4). Statistical significance of the differential response was assessed using a two-sample t-test. Error bars represent the standard error of the mean.

## Supplementary Tables

**Table S1** List of guide RNA sequences and Taqman gene expression primers used for CRISPR/CRISPRi intervention assays

**Table S2** −ΔΔCT values obtained from qPCR studies

**Table S3** List of sample IDs used for this study on the HTAN and GEO portals.

